# Water availability has stronger effects on West Nile virus dynamics in water-limited regions

**DOI:** 10.64898/2026.05.29.728787

**Authors:** Samantha Sambado, Andrew J. MacDonald, Alexandra G. Konings, Erin A. Mordecai

## Abstract

West Nile virus dynamics are shaped by hydrological conditions that influence mosquito habitat and pathogen transmission, but identifying causal relationships is difficult in managed landscapes where irrigation decouples local water conditions from precipitation, complicating climate-disease inference. We address this challenge using a 21-year panel of more than 19 million *Culex tarsalis* mosquitoes from California’s Central Valley, applying fixed-effects panel models to estimate how surface water availability affects mosquito abundance and infection rates while accounting for spatial differences and shared temporal variation. We find that wetter conditions lead to higher mosquito abundance but slightly lower infection rates, suggesting divergent responses of vector population growth and pathogen amplification. These patterns are consistent across multiple hydrological measures, including drought indices, soil moisture, surface water, and evapotranspiration. Effects are strongest in water-limited regions, where hydrological variability is greatest and buffering by snowmelt-fed river systems is weakest. Overall, hydrological conditions exert contrasting effects on key components of West Nile virus dynamics, and these relationships are strongly conditioned by human water management. Our results highlight how irrigation decouples local hydrological conditions from broader climatic variability, underscoring the need for fine-scale hydrological data and panel-based approaches to identify drivers of disease dynamics in human managed landscapes.

## Introduction

West Nile virus, a zoonotic virus that circulates in birds and is transmitted by mosquitoes that lay eggs in standing water, remains a major public health challenge, yet its persistence in semi-arid landscapes presents an ecological paradox (Winters et al. 2008, Cardenas et al. 2011, Godsey et al. 2012, Paull et al. 2017, Snyder et al. 2020, Ward et al. 2023, Sambado et al. 2025, Williamson et al. 2025). In the highest-incidence region of western North America, peak mosquito abundance and WNV transmission occur during the driest months of the year, creating a temporal mismatch between precipitation and disease risk (Reisen et al. 2008a, BARKER et al. 2010). Despite minimal precipitation during the transmission season, WNV continues to circulate, suggesting that local water availability and landscape-level hydrological processes shape transmission in complex ways beyond simple rainfall-mosquito relationships (Reisen et al. 1992, 2008b, Boser et al. 2021, Kovach and Kilpatrick 2024). Such ecological relationships between water management decisions, mosquito abundance, and WNV transmission could in turn present new opportunities for environmental solutions to disease risk.

Previous studies have reported a paradoxical pattern in which an increase in drought severity can reduce mosquito abundance while increasing WNV prevalence (Paull et al. 2017, Sambado et al. 2025). One proposed mechanism is that severe drought reduces the availability of standing water, constraining mosquito development, while also concentrating pathogen-amplifying birds and mosquitoes around remaining water sources, thereby increasing vector-host contact rates (Shaman et al. 2002, 2005, Chase and Knight 2003, Shaman and Day 2005, Skaff and Cheruvelil 2016). Yet, identifying a consistent relationship between drought and mosquito dynamics remains difficult because *Culex* mosquitoes—the primary vectors of WNV—are distributed across the contiguous US and occupy urban, rural, and mixed landscapes that differ substantially in precipitation patterns and thermal suitability (Shocket et al. 2020, Gorris et al. 2021). Precipitation, temperature, and land-use type together shape both natural hydrological conditions and human water-management practices, thereby influencing the presence and persistence of mosquito breeding habitat. Among these vectors, *Culex tarsalis*—the predominant rural vector—is thought to be particularly sensitive to drought conditions because it depends on larger and more persistent aquatic habitat, including wetlands, irrigation ditches, and riparian corridors that are characteristic of rural and agricultural landscapes (Reisen et al. 2009). In contrast, *Culex pipiens* (in the north) and *Culex quinquefasciatus* (in the south) are more strongly associated with urban environments, where they readily exploit small, artificial water sources such as containers, storm drains, and discarded trash (LaDeau et al. 2013, Yitbarek et al. 2023). Consequently, drought may differentially affect vector species depending on their habitat associations and breeding ecology. As a result, the WNV system is highly complex, with multiple vector species, diverse bird hosts, and nonlinear climatic effects, making it difficult to disentangle the many unobserved but impactful drivers of transmission dynamics, particularly related to water (Kilpatrick et al. 2007, DeGroote et al. 2008, Kilpatrick 2011, Harrigan et al. 2014, Rochlin et al. 2019).

To measure water availability on the landscape remote sensing products are commonly used including drought metrics such as the Palmer Drought Severity Index (PDSI), which provide coarse, temporally averaged measures of water availability or landscape “wetness” (Rosile and Bisesi 2017, Paull et al. 2017, Smith et al. 2020, McMillan et al. 2025, Sambado et al. 2025). Mosquitoes and bird hosts, however, can respond to shorter-term and spatially heterogeneous water features (Chase and Knight 2003, Skaff and Cheruvelil 2016, MacDonald et al. 2025). Water availability can take many forms relevant to disease transmission. For example, soil moisture can create hyper-local pools for egg laying and larval development (Ukawuba and Shaman 2018, Davis et al. 2018, Hess et al. 2018). Once mosquitoes emerge as adults they must avoid desiccation and depend on air moisture for their survival (Chuang et al. 2012a, Chuang and Wimberly 2012, Kala et al. 2017, Ward et al. 2023). If mosquitoes survive long enough to obtain a blood meal, relatively larger bodies of standing water, such as irrigation ditches or wetlands, can serve as aggregation sites for mosquitoes and birds, facilitating pathogen transmission (DeGroote et al. 2008, Eisen et al. 2010, Reisen et al. 2013, Kovach and Kilpatrick 2018, 2024, MacDonald et al. 2025). Therefore, multiple potential causal pathways link hydrology, vector ecology, and disease transmission, an increasingly important challenge in an era of precipitation volatility (Swain et al. 2018). Historically, however, testing these relationships across broad spatial and temporal scales was difficult because hydrological observations were often limited to localized measurements, such as rain gauges or field surveys, that could not be easily standardized or integrated with ecological datasets due to spatial or temporal mismatches. Recent advances in satellite remote sensing and the availability of long-term gridded environmental datasets now make it possible to quantify hydrological variability continuously across spatial scales relevant for policy applications. These remotely sensed products, when paired with appropriate assumptions, validation strategies, and robustness checks, can substantially improve the ability to link hydrological variability with long-term ecological dynamics (Proctor et al. 2023, Van Cleemput et al. 2025). Integrating these datasets offers a promising path toward more mechanistic and spatially explicit studies of mosquito ecology and vector-borne disease transmission.

California’s Central Valley provides an ideal system for examining variation in water availability on WNV transmission (Polade et al. 2017). The region is one of the world’s most productive agricultural areas, generating $50 billion annually, but most agricultural production occurs during periods of minimal precipitation (Faunt and Geological Survey (U.S.) 2009). To support crop production, water is intensively managed, accounting for roughly one-sixth of all irrigation water used in the US (Faunt and Geological Survey (U.S.) 2009). The abundance of surface water during warm summer months supports large mosquito populations and persistent WNV transmission since its introduction in 2003 (Reisen et al. 2009, Barker et al. 2010, Snyder et al. 2020, Kovach and Kilpatrick 2024). In this work, we ask: how does water management mediate the effects of drought on WNV risk in the Central Valley? We leverage within-estimator panel models and remotely sensed hydrological variables to approximate causal effects of water availability on WNV risk. We first evaluate broad-scale drought (PDSI) using unit-level (i.e., trapping cluster) and time (year and month) fixed-effects, to control for unobserved, roughly time-invariant characteristics of trapping locations as well as seasonal trends and other time shocks, and examine spatial heterogeneity of drought effects by hydrological regions. We then move beyond a single drought index to investigate more mechanistically-informed hydrological variables, including soil moisture, evapotranspiration, and standing surface water (including irrigation and wetlands), to explore how different dimensions of water availability influence WNV transmission. By combining causal models with hypothesis driven variable selection, we aim to clarify how different dimensions of water availability drive disease risk in a human managed, semi-arid landscape.

## Methods

We investigate how hydrological conditions affect mosquito abundance and WNV transmission by combining long-term mosquito surveillance data with remotely sensed environmental variables in panel regression models. Focusing on California’s Central Valley—a semi-arid, highly irrigated region, where precipitation does not coincide with peak mosquito season—we exploit spatial and temporal variation in surface water availability as a quasi-experimental setting to approximate semi-causal effects (Barker et al. 2010, Hartley et al. 2012). Below, we describe (a) the study system, (b) mosquito surveillance and spatial unit construction, (c) environmental variables, and (d) panel model construction and evaluation.

### (a) Study system

California’s Central Valley covers approximately 18,000 square miles and supplies 40% of the nation’s fruits, nuts and other table foods (Faunt and Geological Survey (U.S.) 2009). The region has a Mediterranean climate, with mild winters and hot summers, with most agricultural production occurring during the summer and fall when precipitation is minimal but temperature and light availability is favorable for crop growth (Fig. 1) (Polade et al. 2017). Rather than being hydrologically uniform, the Central Valley is structured into three major hydrological regions–the Sacramento River, San Joaquin River, and Tulare Basin–that form a gradient in baseline water availability shaped by precipitation, snowmelt-fed river networks, and reliance on managed water infrastructure (Supplementary Fig. 1) (Faunt and Geological Survey (U.S.) 2009). The Sacramento River region in the north is comparatively water-rich, receiving higher precipitation and supported by a dense river network that supplies two-thirds of surface water to the Central Valley, maintaining relatively stable water availability even during drought periods. The San Joaquin River region is intermediate in water availability, with more limited natural inflows and greater dependence on water transfers from the Delta, alongside substantial conversion of historical wetlands to agricultural and urban land uses. In contrast, the Tulare Basin in the south is the most water-limited with low precipitation, strong dependence on the timing and magnitude of Sierra Nevada snowmelt, and heavy reliance on irrigation infrastructure to sustain intensive agriculture. These regional differences reflect the interaction between natural hydroclimatic variability and extensive water management systems that redistribute surface water to meet agricultural, urban, and ecological needs across California. Partly due to this water management system and the region’s long warm season, mosquito-borne diseases are a persistent feature of the Central Valley landscape; since 2003, California has accounted for approximately 18% of all reported neuroinvasive WNV cases in the US (ArboNET) (Reisen et al. 2008a, Snyder et al. 2020, Kovach and Kilpatrick 2024).

**Figure 1.**
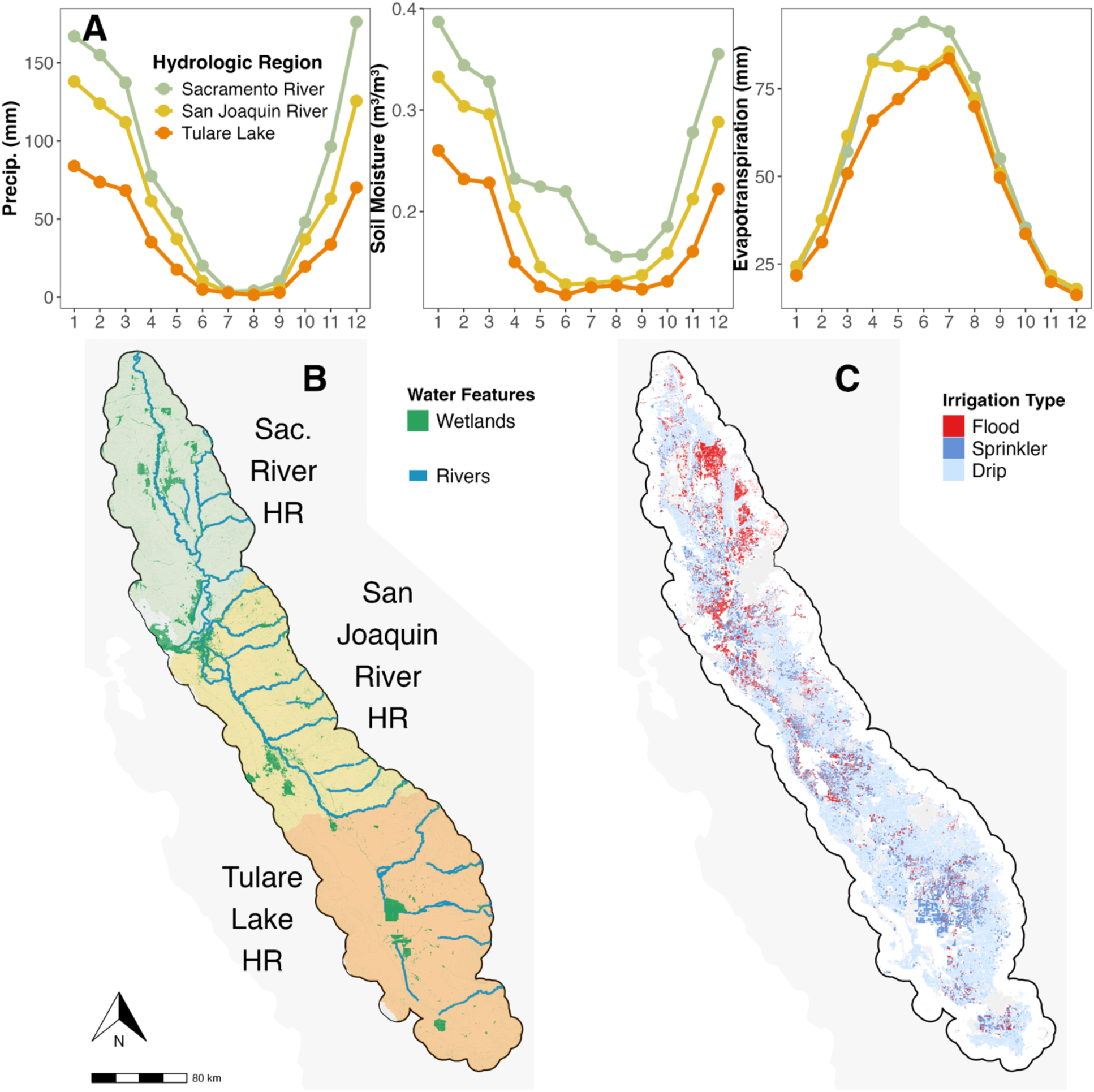
Hydrological context of California’s Central Valley. Spatial and temporal variation in water availability in a semi-arid, intensively managed landscape. (a) Monthly variability of hydrological indicators (precipitation, soil moisture, evapotranspiration), showing spatial gradients in baseline climatic water input. (b) Major rivers and wetlands representing persistent natural aquatic habitat (CDWR). (c) Agricultural land by irrigation type, depicting anthropogenic redistribution of surface water (Land IQ). Spatial boundaries from the ‘tigiris’ package (Walker 2015); alluvial basin from CDWR.

### (b) Mosquito surveillance data

Data were obtained from the CalSurv Gateway through data requests #000092 and #000098 approved by the California Vectorborne Disease Surveillance System, covering 17 counties in California’s Central Valley (Butte, Colusa, Fresno, Glenn, Kern, Kings, Madera, Merced, Placer, Sacramento, San Joaquin, Shasta, Sutter, Tehama, Tulare, Yolo, and Yuba). Data were restricted to April-October, representing peak mosquito activity (Barker et al. 2010, Hartley et al. 2012). Analyses focused on adult female *Cx. tarsalis*, the primary rural mosquito vector in California whose abundance is closely linked with standing water created from agricultural irrigation and natural bodies of water, including rivers and wetlands. Abundance was measured as the average number of mosquitoes per trap night (2003-2023), and WNV infection as the mean minimum infection rate per testing pool (WNV MIR; 2010-2023) (Fig. 2). Additional data filtering and processing details are provided in Supplementary Text 1 and Supplementary Table 1.

**Figure 2.**
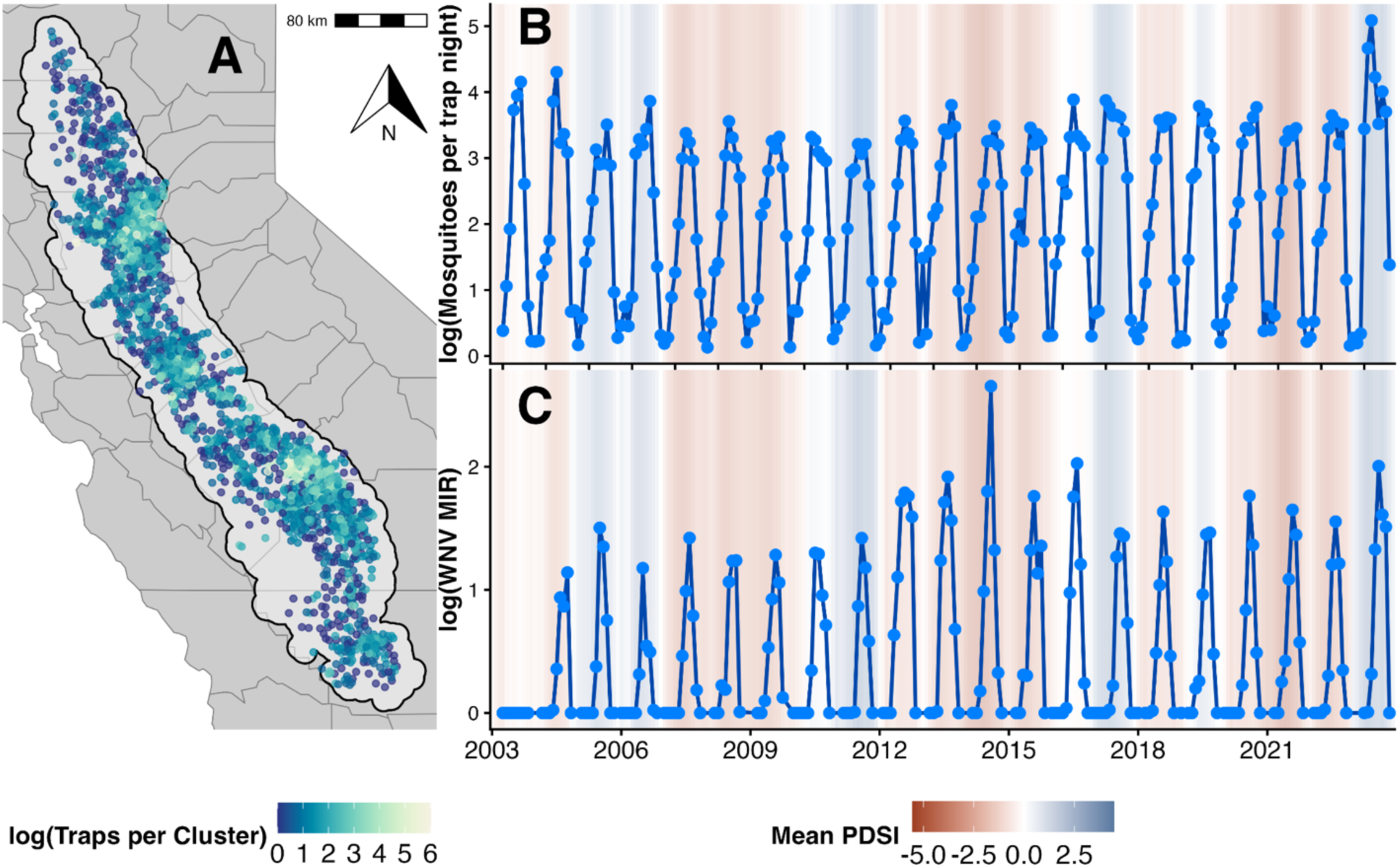
Surveillance data for adult female *Culex tarsalis* mosquito abundance and WNV infection rates. (a) Spatial distribution of sampling clusters, defined as 1.5 km buffers around unique trap stations. (b) Mean mosquito abundance per trap-night and (c) West Nile virus minimum infection rate (WNV MIR) by cluster. Colored boxes show the variation in regional monthly Palmer Drought Severity Index (PDSI; negative to positive; dry to wet), representing landscape wetness conditions.

To account for uneven sampling and spatial autocorrelation due to the non-independence of nearby trap station observations, surveillance trap stations were aggregated into spatial clusters based on estimated dispersal distances of *Culex* mosquitoes (Reisen and Lothrop 1995) (Supplementary Fig. 2). Within each county, we applied complete-linkage hierarchical clustering using the hclust() function (R Core Team, 2024) and generated 1.5 km radius buffers around cluster centroids (MacDonald et al. 2024). Each cluster contained unique trap stations to avoid duplication, and these buffered clusters served as the spatial units for extracting environmental covariates (Supplementary Fig. 3). All outcomes and environmental variables were summarized by cluster-year-month.

### (c) Environmental variables

#### Hydrological treatment

We define each of four hydrological variables as treatments, each capturing a distinct dimension of water availability relevant to mosquito ecology (Table 1). Detailed data sources and processing steps are provided in Supplementary Text 2. First, we include the Palmer Drought Severity Index (PDSI) as an integrated measure of long-term landscape wetness, where lower PDSI values indicate drier conditions and have been associated with reduced mosquito abundance but elevated WNV transmission (Abatzoglou 2013, Sambado et al. 2025) (*data type*: gridded climate dataset; *unit*: index; *spatial resolution*: ∼ 4 km resolution; *source:* PRISM). To isolate specific hydrological mechanisms, we additionally consider three complementary variables. Surface soil moisture represents near-surface water availability and potential breeding habitat for ovipositing mosquitoes (microwave satellite observations, m^3/^m^3^, ∼ 25 km, CCI v09.2) (Gruber et al. 2016, Dorigo et al. 2017, Preimesberger et al. 2021). Standing surface water captures persistent water bodies at landscape scales, reflecting broader ecological conditions that may influence both vector and host communities (satellite imagery, % water-covered pixels within 5 km buffer, 30 m pixels, GSW JRC v1.4) (Pekel et al. 2016). Finally, evapotranspiration (ET) represents the flux of water from land to atmosphere and serves as a proxy for vegetation and atmospheric moisture, which may reduce desiccation risk for adult mosquitoes (satellite and reanalysis model, mm/month, ∼ 10 km, GLEAM v4.2b) (Miralles et al. 2025). These treatments allow us to disentangle pathways through which water availability influences mosquito abundance and infection rates.

**Table 1.** Hydrological pathways linking remotely sensed variables to mosquito ecology. Each row defines a hypothesized causal pathway linking hydrological conditions to *Culex tarsalis* ecology. Pathways connect ecological processes, their corresponding hydrological signals, remotely sensed proxies, and the expected direction of association in within-estimator panel models. These pathways specify the assumed mechanistic structure underlying the analysis. Standing surface water is measured as the percentage of pixels covered by water within a 5 km buffer.

| Ecological process | Hydrological signal | Remotely sensed variable | Hypothesized effect |
| --- | --- | --- | --- |
| **Oviposition habitat availability** | Near-surface, ephemeral water availability | Soil moisture | Higher soil moisture increases availability of oviposition sites, increasing mosquito abundance; effects may be nonlinear under extreme wet conditions due to habitat flushing. |
| **Adult mosquito survival** | Land-atmosphere water flux and vegetation activity | Actual Evapotranspiration | Higher evapotranspiration reflects increased atmospheric moisture and vegetation activity, reducing desiccation stress and increasing adult mosquito survival and abundance. |
| **Host-vector contact rates** | Persistent surface water | Standing surface water | Greater surface water persistence increases aquatic habitat for mosquitoes and birds, redistributing both across water bodies. This may reduce host-vector contact rates and dampen landscape-scale transmission while maintaining local mosquito abundances. |

#### Ecological controls

To better isolate causal effects, we include key time-varying control variables for mosquito physiology and pathogen amplification. Temperature is represented by mean near-surface air temperature (gridded climate dataset, °C, ∼ 1 km, CHELSA) (Karger et al. 2017), included as a quadratic term to capture the nonlinear temperature-abundance relationship (Shocket et al. 2020). Variation in bird host competence is captured using a bird community competence index, which incorporates species-specific infectiousness and seasonal migratory dynamics (Kilpatrick et al. 2007, Hort et al. 2023, MacDonald et al. 2024). Additional details on data processing can be found in Supplementary Text 3.

### (d) Panel models to approximate semi-causal relationships

Vector-borne disease systems, such as WNV, are inherently complex (Fox 1991, Eisenberg et al. 2007, Plowright et al. 2008). Transmission depends on interactions among multiple mosquito vectors, diverse avian hosts, and dynamic environmental conditions (Kilpatrick 2011). Despite this complexity, decades of empirical research have identified relationships—such as how mosquito life history nonlinearly responds to temperature (Shocket et al. 2020) or how certain amounts of water availability can lead to increased abundances (MacDonald et al. 2025)—providing a strong foundation for hypothesis generation (see studies listed in Supplementary Table 2) which should guide causal model formation (Grace and Irvine 2020). Observational studies, however, often struggle to disentangle the effects of hydrology from confounding factors that are multidimensional and highly variable. Instead, causal inference studies have leveraged quasi-random variation in hypothesized environmental drivers to test and quantify their impacts on infectious disease outcomes, controlling for potentially confounding observed and unobserved variation (see examples Dudney et al. 2021, Sambado et al. 2025, Childs et al. 2025). To improve upon these insights, we use within-estimator (fixed-effects) panel models to control for unobserved, time-invariant heterogeneity (Larsen et al. 2019, Byrnes and Dee 2025). By treating each spatial cluster as its own control, these models allow us to isolate the influence of quasi-random temporal fluctuations in hydrological conditions (treatments) on mosquito abundance and WNV infection rates (outcomes), providing a more robust approximation of causal effects in a complex ecological system (MacDonald et al. 2025, Sambado et al. 2025). Our analysis involves two parts.

### Part 1. Overall effect of drought

We first estimated the overall effect of drought, measured by PDSI as a proxy for landscape “wetness”, on *Cx. tarsalis* abundance and WNV infection using the following baseline panel regression:

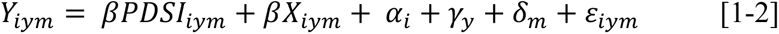

where *i* is a cluster unit, *y* is year, *m* is month. Here, *Y_iym_* represents either mosquito abundance (average mosquitoes per trap night) or WNV MIR (minimum infection rate per testing pool). Both outcomes were log-transformed as log(Y+1) to reduce right-skewed distributions characterized by many low values and a small number of high observations. *X_iym_* includes outcome-specific covariates. For both models, we included standardized mean temperature and its quadratic term to capture nonlinear thermal effects. For WNV infection models, we additionally included a standardized index of bird community competence. These models include a cluster unit (*i*) fixed effect to control for time-invariant spatial heterogeneity, along with year (*γ_y_*) and month (*δ_m_*) fixed effects to control for interannual variation and seasonality. *ε_iym_* is the error term. Robustness checks included alternative specifications, including fixed-effect specifications, model types (Poisson vs negative binomial), lagged treatment effects (1- to 5-month lag), covariate functional forms, and varying spatial cutoffs for Conley standard errors that account for spatial autocorrelation (Supplementary Text 4, 5).

To explore spatial heterogeneity in the mosquito-PDSI relationship, we extended the baseline model by interacting PDSI with region indicators:

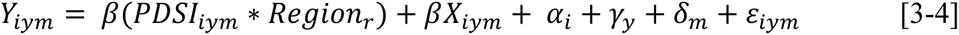

where *Region_r_* denotes hydrological regions (Sacramento River, San Joaquin River, Tulare Lake). This specification allows the marginal effect of drought to vary across regions, with the hypothesis that drought effects on mosquito outcomes are strongest in more water-limited regions such as the Tulare Basin, and weakest in the comparatively surface water-rich Sacramento River region. As an additional robustness check, we estimated models allowing the effect of PDSI to vary at finer spatial scales by interacting PDSI with county indicators (Supplementary Text 6).

### Part 2. Mechanistic links between hydrological variables and mosquito ecology

To identify which components of water availability are most strongly associated with mosquito abundance and WNV transmission, we re-estimated the fixed-effects panel models using alternative hydrological indicators in place of PDSI. These included standardized measures of soil moisture, evapotranspiration, and standing surface water (within 5 km radius of a cluster unit). The specification was:

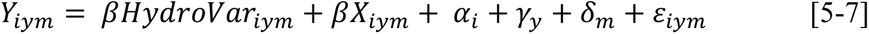

Where *HydroVar_iym_* represents each hydrological variable: soil moisture as a proxy for near-surface water availability, evapotranspiration as a proxy for atmospheric water flux, and standing surface water. Because these hydrological indicators capture distinct but related processes, they were not included simultaneously in the same model. To assess temporal dynamics, we estimated models with 0-, 1-, 2-, and 3-month lags for each hydrological variable, where the hydrological predictors preceded the response variable by the specified number of months and a 0-month lag represents contemporaneous conditions (Supplementary Text 7). All other terms are defined in the baseline model (Equations 1-2). This approach isolates the association between specific hydrological processes and mosquito outcomes, while controlling for time-invariant spatial characteristics and shared temporal trends.

All models used Conley standard errors to account for spatially correlated residuals. For estimation, we used ordinary least squares regression with the ‘fixest’ package (Bergè 2019). All covariates were standardized for comparability, and variance inflation factors were checked to avoid collinearity. Additional sensitivity analyses addressed potential threats to causal inference, including measurement error, spatial scale dependence, and external validity (Ferraro 2009, Siegel and Dee 2025, Dudney et al. 2025). Full details are provided in Supplementary Text 4, 5, 6, 7, 8.

## Results

### Surveillance data and mosquito infection rates

The dataset included 19,440,201 adult female *Cx. tarsalis* mosquitoes collected across 1,214,538 trap nights. A total of 24,372 unique trap stations were aggregated into 2,818 spatial clusters. Virus testing included 414,722 mosquito pools, representing 13,705,162 individual mosquitoes tested (Table 2, Supplementary Table 3). Overall, 4.1% (16,811/414,722) of mosquito pools tested positive for West Nile virus (Fig. 2, Supplementary Fig. 4, 5).

**Table 2.** Summary statistics for *Culex tarsalis* surveillance data (April-October, 2003-2023). Counties are grouped by hydrological region: Sacramento River (SAC), San Joaquin River (SAN), and Tulare Lake (TUL). Sampling effort is summarized as the total number of clusters and trap nights, along with cumulative adult female mosquito counts. County-level vector and environmental conditions are reported as mean (standard deviation): West Nile virus minimum infection rate (WNV MIR) and Palmer Drought Severity Index (PDSI). Human West Nile virus incidence is provided as contextual information, expressed as mean annual incidence per 100,000 population over the study period, with total case counts in parentheses, based on ArboNET surveillance data.

| County | Clusters | Trap Nights | Mosquitoes | WNV MIR | PDSI | Human Incidence |
| --- | --- | --- | --- | --- | --- | --- |
| **Butte**<br>(SAC) | 43 | 14,497 | 651,748 | 2.67 (5.8) | -1.22 (2.9) | 5.99 (281) |
| **Colusa**<br>(SAC) | 7 | 2,128 | 106,465 | 2.24 (6.0) | -1.30 (3.0) | 5.10 (23) |
| **Fresno**<br>(SAN) | 412 | 14,050 | 1,096,600 | 3.10 (9.7) | -0.99 (3.0) | 1.91 (398) |
| **Glenn**<br>(SAC) | 13 | 5,125 | 361,359 | 2.76 (4.8) | -1.36 (3.1) | 16.97 (100) |
| **Kern**<br>(TUL) | 181 | 21,395 | 2,395,117 | 3.54 (10.3) | -0.71 (2.9) | 3.11 (583) |
| **Kings**<br>(TUL) | 70 | 5,374 | 699,184 | 5.84 (14.0) | -0.93 (2.9) | 3.06 (97) |
| **Madera**<br>(SAN) | 108 | 13,844 | 452,135 | 6.46 (12.4) | -0.96 (3.1) | 2.41 (79) |
| **Merced**<br>(SAN) | 109 | 24,410 | 514,133 | 2.99 (16.8) | -1.07 (3.0) | 1.93 (111) |
| **Placer**<br>(SAC) | 130 | 9,252 | 596,310 | 1.90 (13.6) | -0.81 (2.7) | 1.35 (111) |
| **Sacramento**<br>(SAC) | 264 | 88,418 | 1,074,754 | 4.82 (33.1) | -1.06 (2.9) | 1.27 (410) |
| **San Joaquin**<br>(SAN) | 124 | 12,797 | 976,250 | 4.20 (22.4) | -1.28 (2.9) | 1.25 (197) |
| **Stanislaus**<br>(SAN) | 176 | 12,733 | 712,704 | 2.22 (23.2) | -1.07 (2.8) | 3.76 (431) |
| **Sutter**<br>(SAC) | 33 | 11,951 | 891,833 | 3.61 (8.65) | -1.27 (3.0) | 3.86 (78) |
| **Tehama**<br>(SAC) | 13 | 2,103 | 21,650 | 2.13 (4.5) | -1.15 (2.8) | 4.53 (61) |
| **Tulare**<br>(TUL) | 198 | 15,449 | 330,804 | 4.43 (13.6) | -0.97 (3.1) | 2.69 (262) |
| **Yolo**<br>(SAC) | 102 | 64,146 | 2,561,061 | 1.78 (8.9) | -1.18 (3.1) | 3.61 (166) |
| **Yuba**<br>(SAC) | 17 | 10,355 | 316,028 | 2.86 (5.22) | -1.20 (2.9) | 4.18 (68) |

### Divergent effects of landscape wetness on mosquito abundance and infection rates

Higher PDSI (greater landscape wetness) was strongly associated with increased mosquito abundance but slightly reduced WNV MIR (Fig. 3A). These results were supported by extensive robustness checks across alternative model structures, lag specifications, functional forms, distributional assumptions, spatial error corrections, vector species, and geographic subsamples. For abundance, a one-unit increase in PDSI was associated with a 16.5% increase in mosquito abundance (Beta = 0.15, SE = 0.017, 95% CI [0.12, 0.19], p < 0.001). A one-unit increase in PDSI reflects a meaningful shift in overall landscape moisture conditions rather than short-term weather variability. For context, the range of PDSI over the entire 21-year study period was -8.79 to 12.92, indicating substantial variation in long-term landscape moisture conditions. Temperature showed a nonlinear effect, with abundance peaking near 29 °C (Supplementary Text 4). For infection, PDSI was negatively associated with WNV MIR (Beta = -0.053, SE = 0.014, 95% CI [-0.08, -0.03], p < 0.001), corresponding to a 5.4% decrease in infection rate per unit increase in PDSI (Fig. 3A). WNV infection rates showed no significant linear association with temperature, but a significant quadratic term suggests evidence of nonlinearity, with infection at warmer temperatures (Supplementary Text 5). Bird community competence was positively associated with WNV infection rates in pooled OLS models, but this association became negative and was not statistically significant in models including cluster, year, and month fixed effects.

**Figure 3.**
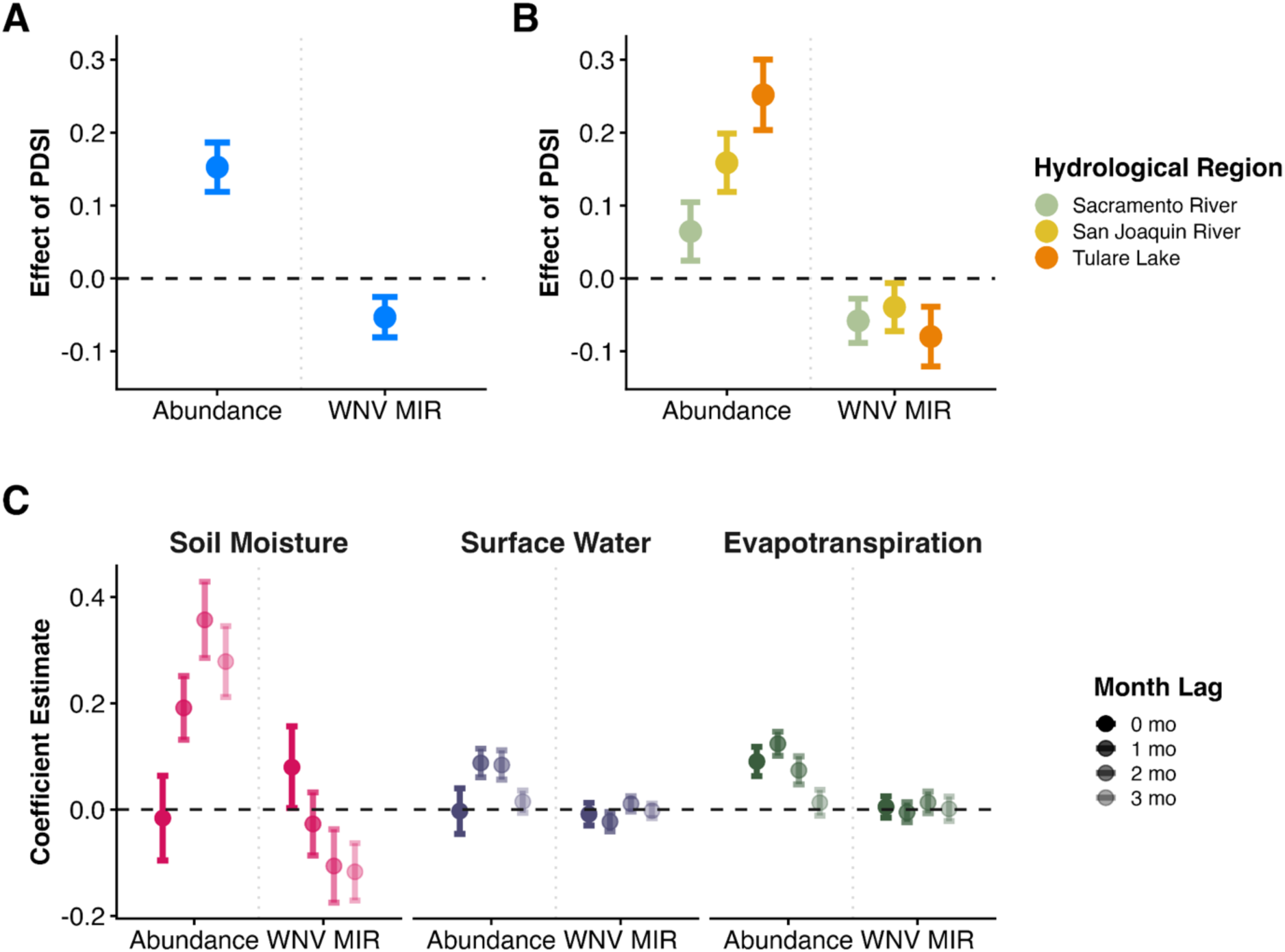
Estimated effects of hydrological variability on mosquito abundance and infection rates from within-estimator panel models. Point estimates represent regression coefficients; error bars denote 95% confidence intervals. The primary exposure of interest is Palmer Drought Severity Index (PDSI; top row), with estimates supported by robustness and sensitivity analyses (see SI). (a) Average effect of PDSI on mosquito outcomes. (b) Spatial heterogeneity in PDSI effects across hydrological regions. (c) Effects of mechanistic hydrological variables (soil moisture, surface standing water within a 5 km buffer, and actual evapotranspiration) on mosquito outcomes across temporal lags (0-3 months).

### Drought effects vary spatially across hydrological regions

The relationship between PDSI and mosquito outcomes were similar in direction to our baseline model, but the magnitude of effect differed by hydrological region (HR; Fig. 3B). For abundance, all regions showed positive associations, with the strongest effect in Tulare Lake HR (Beta = 0.25, SE = 0.025, p < 0.001), the southernmost region of the Central Valley, followed by San Joaquin River HR (Beta = 0.16, SE = 0.020, p < 0.001) and Sacramento River HR (Beta = 0.064, SE = 0.020, p < 0.001; Fig. 3B). For WNV MIR, associations were negative across all regions, with Tulare Lake HR (Beta = -0.080, SE = 0.021, p < 0.001) again having the strongest effect, followed by Sacramento River HR (Beta = -0.058, SE = 0.015, p < 0.001), and San Joaquin River HR (Beta = -0.039, SE = 0.017, p = 0.002; Fig. 3B). County-level analyses broadly mirrored these patterns, although some counties exhibited substantial uncertainty, particularly those with smaller sample sizes or sparse infection data, as indicated by some confidence intervals overlapping zero (Supplemental Text 6). Overall, wetter conditions strongly increased mosquito abundance but slightly decreased infection rates, with the strongest responses observed in more water-limited regions. This pattern was most pronounced in the Tulare Basin, where low precipitation and singular major river system - much of whose flow is heavily diverted for agricultural use - provide limited buffering against drought-driven shifts in mosquito dynamics, in contrast to the Sacramento River region, which is sustained by a dense network of large, perennial rivers that deliver substantial surface water inputs and buffer against precipitation variability.

### Hydrological drivers differentially impact mosquito outcomes

Hydrological effects on mosquito outcomes were time-dependent and differed between abundance and infection processes (Supplementary Text 7; Fig. 3C). For abundance, in general, more available water resulted in significant increases. At the contemporaneous scale, higher soil moisture was associated with reduced abundances, but this relationship reversed at 1-3 month lags (Beta range: 0.19 - 0.29). This suggests short-term suppression followed by delayed habitat-driven increases. Standing surface water showed similar but weaker patterns (Beta range: 0.01 - 0.12). Evapotranspiration was positively associated with abundance at 0-3 month lags, declining over time. For WNV infection, associations were weaker and temporally segmented into early (0-1 month) and delayed (2-3 month) regimes. Soil moisture initially increased infection rates (Beta = 0.08, SE = 0.039, p < 0.05), but then declined over time with significant associations during 2-3 month lags (Beta range: -0.11 - -0.12). Standing surface water showed an initial negative response, followed by a short-lag positive response, that declined at longer lags, but these were very weak relationships. Evapotranspiration was never significantly associated with infection rates, unlike abundances. Of all the hydrological predictors, soil moisture exhibited the strongest relationships, increasing abundance while decreasing infection at lags of 1-3 and 2-3 months, respectively.

## Discussion

Across California’s Central Valley, wetter hydrological conditions were associated with higher *Cx. tarsalis* abundance but lower WNV infection rates, indicating that vector population dynamics and pathogen transmission respond differently to variation in water availability. These relationships were spatially heterogeneous, with the strongest effects observed in the southern Central Valley, where precipitation is minimal, river networks are limited and highly dependent on variable Sierra Nevada snowpack, and irrigation for agriculture constitutes a major source of surface water across the landscape. Directional effects were broadly consistent across remotely sensed hydrological variables and temporal lags, aligning with expectations regarding mosquito habitat formation, population growth, and viral amplification (Reisen et al. 2009, Barker et al. 2010, Smith et al. 2020, Sambado et al. 2025). Our results suggest that climate-disease relationships are mediated by human water management, which reshapes how hydrological variability translates into ecological and epidemiological risk.

The effects of drought were directionally consistent across hydrological regions but differed substantially in magnitude, with the strongest responses observed in the southern Tulare Basin. This spatial heterogeneity likely reflects regional differences in baseline hydrology and water management (Reisen et al. 2008a, Faunt and Geological Survey (U.S.) 2009, Barker et al. 2010, Boser et al. 2024). The northern Central Valley receives substantially greater winter precipitation and streamflow than southern regions, supporting extensive river networks fed by the Klamath Mountains, Sierra Nevada, and Coastal Ranges. As a result, the Sacramento River region maintains relatively stable surface water availability, even during drought years, through large and persistent snowmelt-fed river systems. In contrast, the southern Central Valley relies more heavily on hydrologically variable Sierra Nevada snowpack, particularly through the Kern River system, making water availability more sensitive to interannual drought conditions. The San Joaquin River region is intermediate, historically supported by extensive Delta wetlands that may partially buffer drought impacts but remain vulnerable during prolonged or severe dry periods. These natural hydrological differences are further modified by extensive water infrastructure. The Tulare Basin imports more surface water than any other region in California through canals and irrigation networks that sustain intensive agriculture despite chronically low winter precipitation. At the same time, irrigation practices differ regionally: the Sacramento River region contains extensive flooded rice agriculture that maintains persistent aquatic habitat, whereas agriculture in the Tulare Basin more commonly relies on drip and sprinkler irrigation systems that generate more spatially and temporally variable mosquito habitat. Consequently, broad drought metrics such as PDSI may represent only one mosquito-relevant water availability metric in managed agricultural systems where irrigation redistributes water independently of local precipitation (Winters et al. 2008, Eisen et al. 2010, Kovach and Kilpatrick 2018, 2024, Bhattachan et al. 2021, Wheeler et al. 2022, Williamson et al. 2025). Our findings suggest that human-managed water systems can either buffer or amplify ecological responses to drought, thereby limiting the explanatory power of climatic drought indices alone (Godsey et al. 2012).

To examine the mechanisms underlying these relationships, we evaluated multiple remotely sensed indicators representing distinct dimensions of water availability, including soil moisture, standing surface water within a 5 km buffer, and evapotranspiration. Across indicators, mosquito abundance and WNV infection rates responded in opposite directions, similar to responses observed for PDSI, yet with some distinctions that aligned with ecological expectations (Shaman et al. 2005, Paull et al. 2017, Sambado et al. 2025). Notably, increases in soil moisture and surface water were associated with short-term declines in abundances, followed by delayed positive associations. These initial declines are consistent with the “flushing out” hypothesis (see conceptual Fig. 5 of Caldwell et al. 2021) in which rapid inundation temporarily disrupts larval habitat before more stable aquatic environments develop (MacDonald et al. 2025). Temporal responses also differed systematically between abundance and infection outcomes. Mosquito abundance responded more directly and linearly to changing habitat suitability, including with evapotranspiration, whereas infection rates exhibited delayed responses consistent with the additional ecological processes required for enzootic amplification (Reisen et al. 2006, 2009). Infections in mosquitoes requires a sequence of events: mosquitoes must first emerge from a water source and locate a blood meal, often from birds aggregated around limited water resources; transmission then depends on feeding upon an infected host, followed by the completion of the virus’s extrinsic incubation period before the mosquito becomes capable of transmitting the virus. Collectively, these lag structures support the interpretation that hydrological variability directly influences WNV transmission through ecologically mediated pathways rather than purely coinciding in time and space.

Previous studies of *Culex* mosquitoes and WNV transmission across North America have consistently identified temperature and precipitation as important environmental drivers, while a smaller but growing literature has emphasized broader hydrological variability, including drought indices, soil moisture, evapotranspiration, and surface water dynamics (Table S2). In semi-arid regions such as Arizona, California, Colorado, and Texas, where temperatures are frequently permissive for WNV transmission, variation in water availability may impose stronger ecological constraints on mosquito populations than temperature alone, unlike more northern states like Illinois, New York, South Dakota, and Ohio (Morin and Comrie 2013, Keyel et al. 2021, Gorris et al. 2023). However, mosquito habitat is shaped not simply by precipitation itself, but by the persistence of standing water and connectivity of water bodies across the landscape (Reisen et al. 1992, Chase and Knight 2003, Barker et al. 2009, Skaff and Cheruvelil 2016, MacDonald et al. 2025). Studies from wetland ecosystems outside of California have shown that hydrological connectivity can strongly influence mosquito abundances and human WNV risk by altering the spatial arrangement of aquatic habitat and persistence of aquatic habitat, as well as the broader ecological trophic cascades that sustain WNV transmission during drought conditions (Chase and Knight 2003, Skaff and Cheruvelil 2016). Similar processes may operate across California’s Central Valley, where regional differences in river networks and wetland extents can shape how water moves and persists across agricultural landscapes. Our findings therefore support the view that hydrological variability is not a single environmental exposure, but rather a set of interacting processes operating across multiple spatial and temporal scales that can differentially affect complex transmission cycles (Chuang et al. 2012b, 2012a, Chuang and Wimberly 2012, 2012, Kala et al. 2017, Ukawuba and Shaman 2018, Hess et al. 2018, Ward et al. 2023). In agricultural regions where precipitation is decoupled from peak mosquito season, soil moisture may provide a more informative measure of hydrological conditions at broader spatial scales because it integrates both recent precipitation and irrigation-driven water availability. Consistent with this interpretation, soil moisture exhibited larger effect sizes than standing water or evapotranspiration in our analyses, although the relative magnitude of these associations should be evaluated across multiple remotely sensed products. Integrating remotely sensed hydrological observations with ecological surveillance data may consequently improve mechanistic understanding of vector-borne disease dynamics in water-limited and highly managed environments.

A key consideration in interpreting these results is that remotely sensed hydrological variables capture ecological water availability only indirectly, as proxies for fine-scale hydrological conditions that shape mosquito habitat. For example, soil moisture is unlikely to influence mosquito abundance directly but instead reflects broader landscape wetness that is associated with the formation and persistence of small, often ephemeral standing water bodies that support larval development. More generally, remotely sensed hydrological indicators simplify a highly heterogenous and dynamic water landscape, introducing two distinct sources of uncertainty when scaling ecological processes. First, intrinsic measurement error arises from biases in retrieval algorithms and model assumptions, even when spatial and temporal scales are perfectly aligned (Proctor et al. 2023, Van Cleemput et al. 2025). Second, representativeness error arises from mismatches in spatial resolution and temporal aggregation between satellite products and the fine-scale ecological processes they are used to approximate, and varies across space (Formanek et al. 2026a, McCormick et al. 2026). In this study, we relied on relatively coarse-resolution products for soil moisture and evapotranspiration due to the data availability over the full study period. This may be particularly consequential for evapotranspiration products like GLEAM, where coarse spatial aggregation could contribute to weaker observed associations. Future work would benefit from higher-resolution datasets such as OpenET for evapotranspiration or SMAP for soil moisture (Boser et al. 2024, Formanek et al. 2026b). Additional limitations of our results arise from the structure of the hydrological datasets themselves. The soil moisture or standing surface water product does not distinguish among ecologically distinct water sources, including wetlands, irrigated agriculture, and riverine systems, despite their potentially different contributions to larval habitat suitability and transmission dynamics. Similarly, monthly aggregations may obscure short-term inundation and drying cycles that are critical to mosquito-life history processes that can occur on the weekly scale. The monthly timestep was dictated by the availability of the standing water product, which is constrained by Landsat revisit frequency and cloud-free observations. This constraint does not apply uniformly across hydrological variables, meaning the degree of temporal aggregation bias varies across exposure variables. Despite these limitations, the consistency of estimated effects across multiple hydrological indicators, as well as across independent soil moisture products with differing spatial resolutions (Supplementary Text 8.2), suggest that the main findings are robust to alternative measurement specifications. However, residual exposure misclassification and unobserved hydrological heterogeneity are likely to persist given the inherent complexity of water dynamics in managed landscapes, underscoring the fundamental challenge of representing hydrological variability in discretized datasets. Addressing this limitation will require continued integration of high-resolution remote sensing, hydrological modeling, and ecological surveillance to better align measurement frameworks with biologically relevant processes and to improve inference in climate-sensitive disease systems.

Finally, our results highlight how human management of water can reshape climate-disease relationships in agricultural landscapes (Hess et al. 1970, DeGroote et al. 2008, Liu et al. 2008, Eisen et al. 2010, Baeza et al. 2011, Kovach and Kilpatrick 2018, 2024, Williamson et al. 2025). Across California’s Central Valley, irrigation redistributes water through extensive canal networks, groundwater extraction, and diverse irrigation systems, partially decoupling mosquito habitat availability from natural precipitation variability. However, irrigation practices differ substantially among crops and regions, producing different ecological consequences for mosquito populations (Boser et al. 2021, 2024). In the northern and north-central Central Valley, flooded rice agriculture creates extensive and persistent aquatic habitat that is strongly associated with increased *Cx. tarsalis* abundance and elevated WNV risk (Garcia et al. 1992, Lawler and Dritz 2006). Mosquito population dynamics in these regions closely track the seasonal timing of rice cultivation and flooding, despite intensive coordination between vector-control agencies and rice growers to balance public health protection with agricultural production (Reeves et al. 1990, Wheeler et al. 2022). By contrast, sprinkler and drip irrigation systems more commonly found in the southern Central Valley generally create smaller and less persistent aquatic habitats. Legislation such as the Sustainable Groundwater Management Act (SGMA) is expected to substantially reshape irrigation practices in California, with projections of more than 500,000 acres of agricultural land being fallowed. These changes are likely to alter landscape-scale water inputs, and in turn, influence habitat availability for mosquitoes and pathogen-amplifying bird hosts. As precipitation becomes less predictable and average temperatures increase, demand for irrigation is expected to increase in many agricultural regions, motivating the need for data-driven approaches that integrate agroecological processes with remote sensing while explicitly accounting for vector-borne disease risk (Boser et al. 2021, 2024). Previous studies have shown that irrigation can substantially increase *Culex* abundance, reduce seasonal variability in mosquito populations, and explain considerable spatial variation in human WNV incidence (Eisen et al. 2010, Cardenas et al. 2011, Kovach and Kilpatrick 2018, 2024). In some regions, and almost always in California, irrigation inputs now exceed precipitation during the summer growing season, effectively buffering mosquito populations from climatic water limitations (DeGroote et al. 2008, Faunt and Geological Survey (U.S.) 2009). More broadly, these findings suggest that engineered water systems do not simply respond to climate variability, but actively mediate ecological transmission pathways. Beyond the *Culex* system, evidence for human-mediated water-disease relationships is growing for schistosomiasis, malaria, dengue, cholera, and *Giardia* to name a few, but largely impactful disease systems, highlighting the broader relevance of hydrological infrastructure for infectious disease dynamics (Lerer and Scudder 1999, Baeza et al. 2011, Lindahl and Grace 2015). As climate change, agricultural intensification, and water scarcity increase reliance on irrigation and other hydrological infrastructure worldwide, integrating long-term disease surveillance, environmental monitoring, and hydrological observations may become increasingly important for anticipating unintended public health consequences of water policy and infrastructure decisions.

## Acknowledgements

We acknowledge Zoe Rennie, who developed the bird community competence layer. We thank the vector control agencies who contributed the data to this project: Alameda County MAD, Arbovirus Field Station, Bakersfield (UC Davis), Antelope Valley MVCD, Butte County MVCD, Calaveras County of Environmental Health Department, Colusa MAD, Contra Costa County MVCD, Delano MAD, Delta VCD, East Side MAD, Fresno MVCD, Fresno Westside MAD, Glenn County MVCD, Kern MVCD, Kings MAD, Lake County VCD, Madera County MVCD, Merced County MAD, Oroville Mosquito Abatement District, Placer MVCD, Sacramento-Yolo MVCD, Santa Clara MVCD, Shasta MVCD, San Joaquin County MVCD, Solano County MAD, Sutter-Yuba MCVD, Tehama County MVCD, Tulare MAD, Turlock MAD, Vector-Borne Disease Section, California Department of Public Health, and West Side MVCD.

## Data materials and availability

All original data sources used in this study are publicly available (except mosquito surveillance data) and are detailed in Supplementary Text 1 & 2, Supplementary Table 1, which include direct links to each dataset. Code used to reproduce all analyses and figures is available on GitHub (https://github.com/sbsambado/wnv_water_ca) and will be archived in a Dryad repository (10.5061/dryad.cnp5hqcm6). The mosquito surveillance data were acquired from the California Vectorborne Disease Surveillance System (CalSurv Gateway; https://ca.vectorsurv.org/) via data requests #000092 (approved on 15 January 2026) and #000098 (approved on 17 March 2026). For figure visualizations, spatial boundary data include California state and county boundaries from TIGER/Line shapefiles using ‘tigris’ package (Walker 2015), and the Central Valley alluvial boundaries from the California Department of Water Resources (CDWR; https://water.ca.gov/). For Figure 1, hydrological features were compiled from the National Hydrography Dataset (NHD) (https://www.usgs.gov/national-hydrography/national-hydrography-dataset) and the National Wetlands Inventory (NWI; https://www.fws.gov/program/national-wetlands-inventory). Irrigation data came from crop mapping products developed by LandIQ (https://www.landiq.com/). Precipitation data (Fig. 1A, 1B) were from PRISM Group, Oregon State University (Abatzoglou 2013). Human neuroinvasive case data came from ArboNET (https://www.cdc.gov/vector-borne-diseases/php/arbonet/).

