## Supplementary Information for "Water availability has stronger effects on West Nile virus dynamics in water-limited regions"

This document contains supplementary figures, tables, texts and references supporting the main manuscript.

*Figures*

**Figure S1.** Map of water management in California
**Figure S2.** Distribution of trap stations per cluster
**Figure S3.** Map of cluster locations
**Figure S4.** Mosquito abundance by county-year
**Figure S5.** Mosquito infection rate by county-year
**Figure S6.** Time series of hydrological variables
**Figure S7.** Bird community competence by county-month

*Tables*

**Table S1.** Data sources and resolutions
**Table S2.** Previous empirical *Culex*–climate studies
**Table S3.** Summary statistics

*Text*

**Text S1.** Mosquito surveillance and data processing
**Text S2.** Hydrological variables - rationale and processing
**Text S3.** Other data justifications
**Text S4.** Main abundance panel model evaluation and robustness checks
**Text S5.** Main WNV MIR panel model evaluation and robustness checks
**Text S6.** Spatial heterogeneity in panel models
**Text S7.** Hydrological variable panel models
**Text S8.** Sensitivity analysis

*References*

**Supplementary Figures**

**Figure S1 Map of water management and hydrological context in California**. (a) Total annual precipitation (mm; 1991-2020 normals (Abatzoglou 2013)), showing spatial gradients in baseline climatic water input. (b) Mean monthly precipitation (mm) by hydrological region, illustrating seasonal decoupling between rainfall and peak mosquito activity. (c) Expands on Figure 1 by showing multiple wetland and river types. Wetland data are from the USGS National Wetlands Inventory, and rivers are classified by 'ftype' codes from the National Hydrography Dataset (NHD).

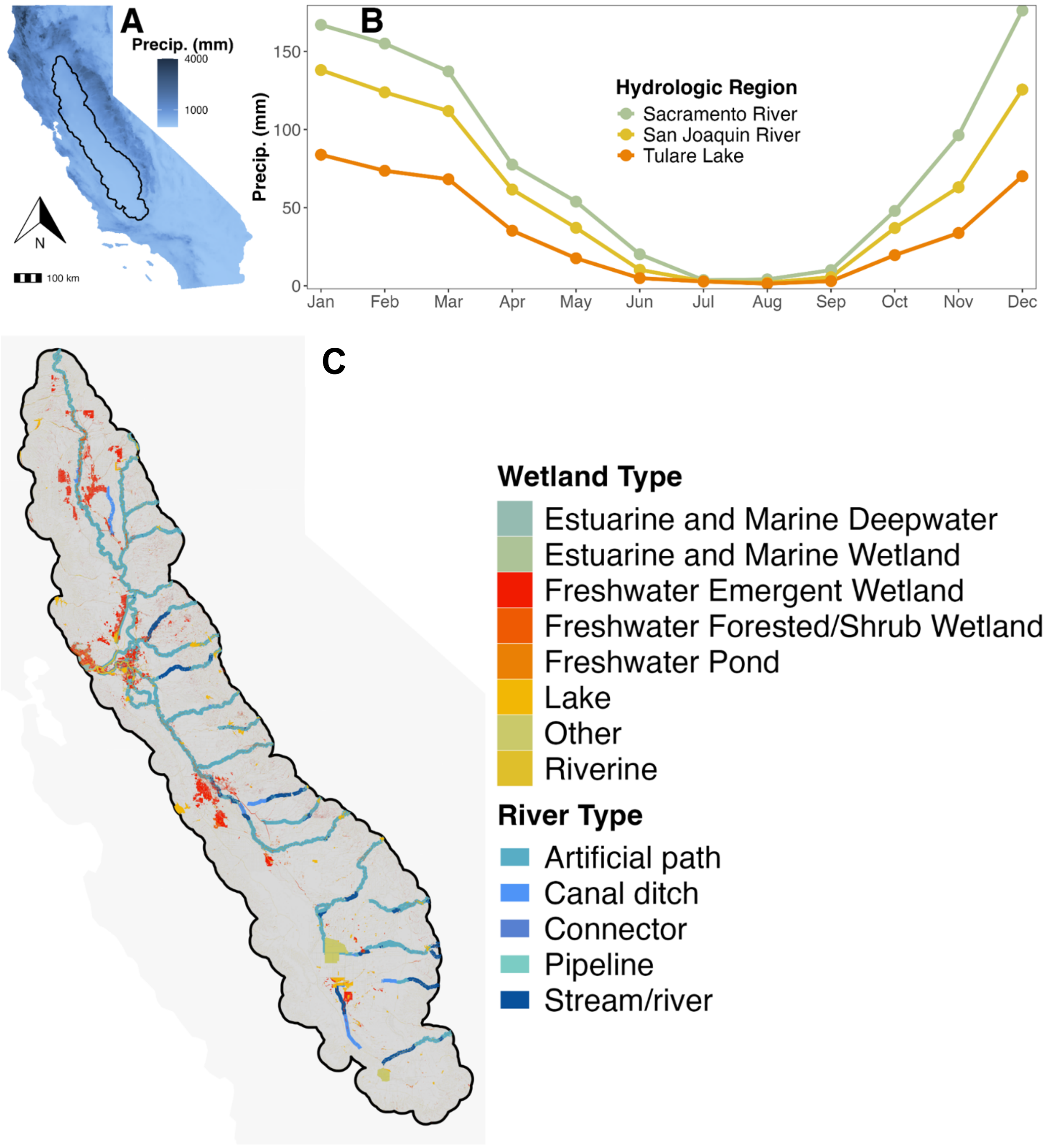

**Figure S2 Distribution of trap stations per cluster.** Histogram of the number of unique trap stations per cluster across the study period, limited to clusters with fewer than 100 stations. Trap stations were established by vector control districts across California.

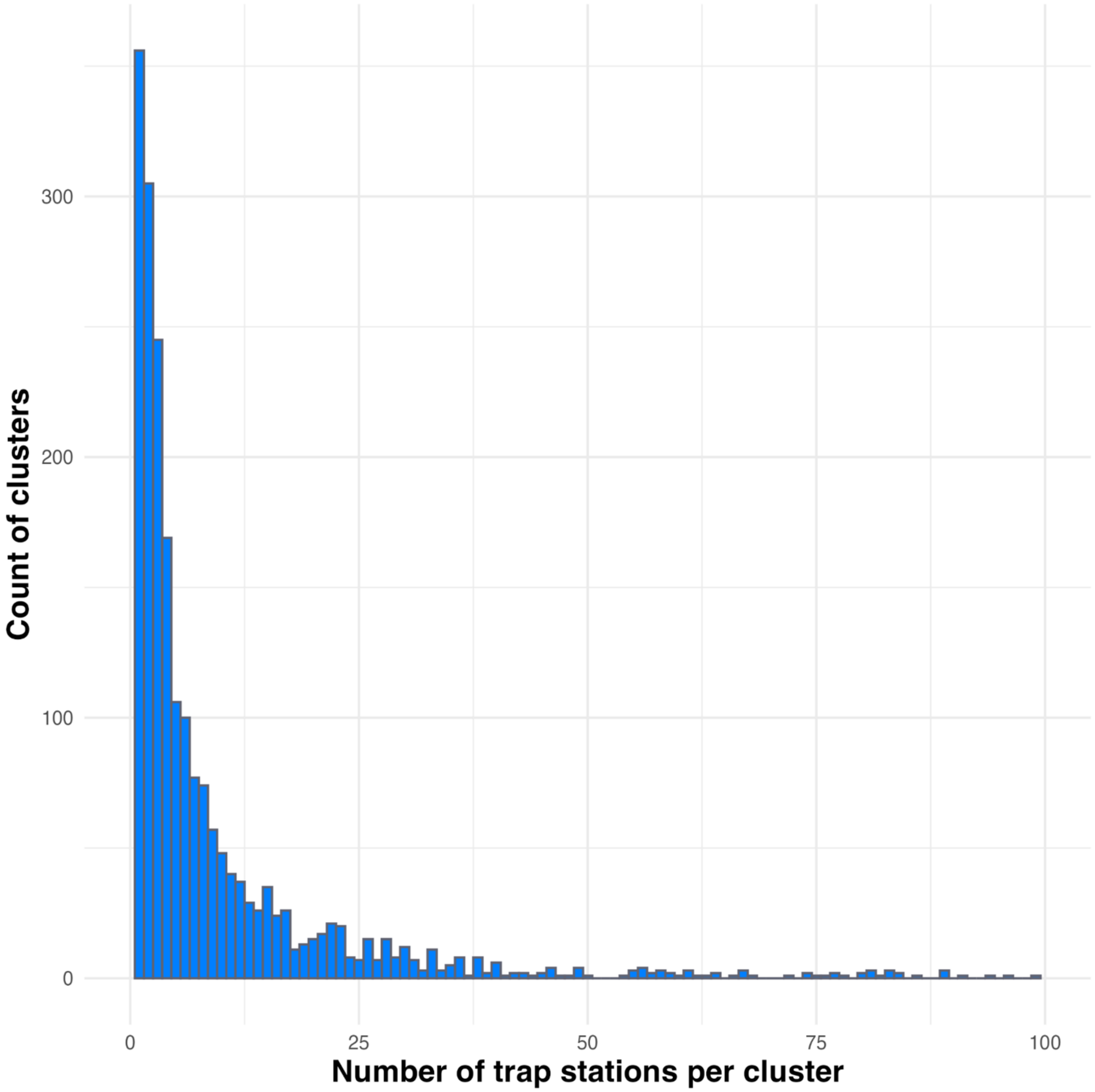

**Figure S3 Map of cluster locations.** Locations of mosquito trap clusters in California's Central Valley. The subplot highlights cluster locations within hydrological regions (HRs). Points are colored by the log-transformed number of traps per cluster. Cities with populations over 100,000 are labeled.

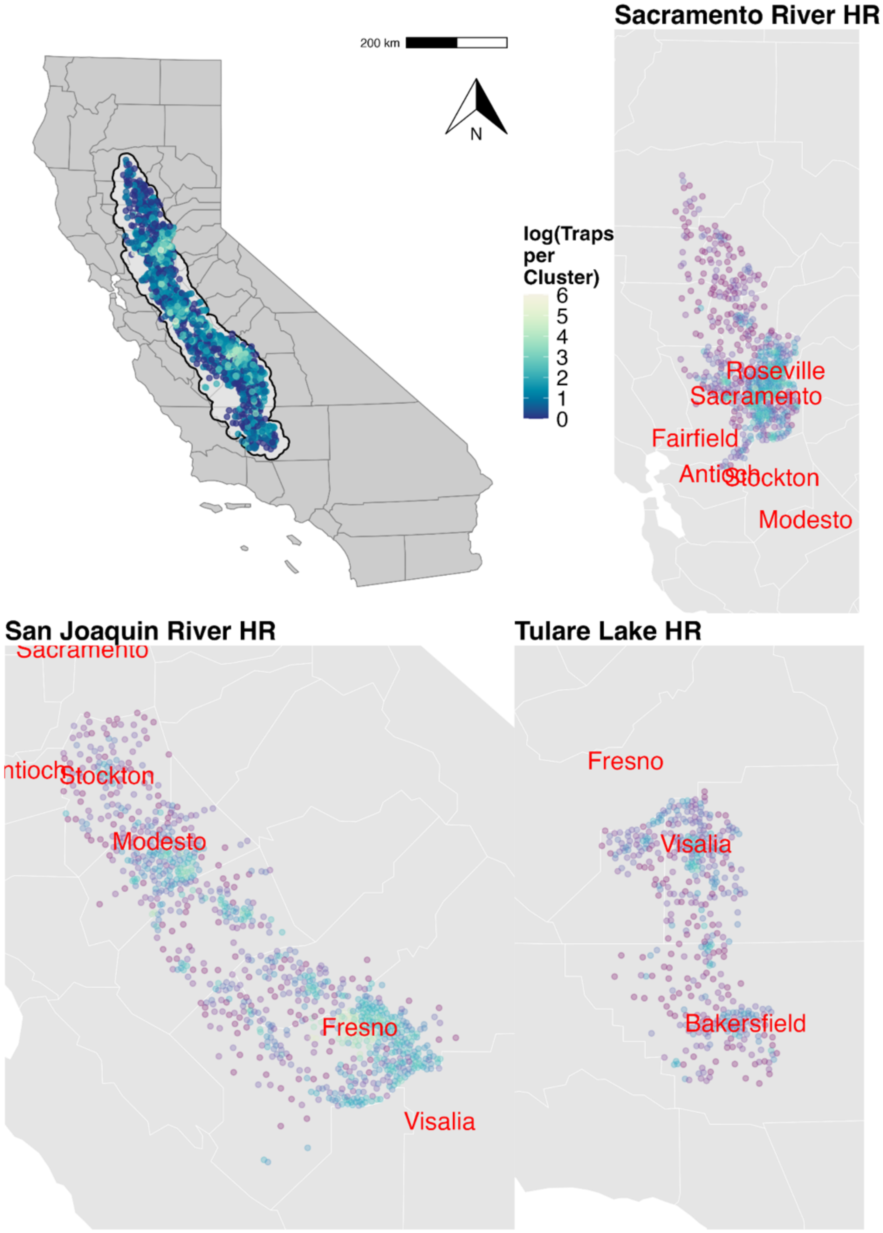

**Figure S4 Mosquito abundance by county-year.** Data from CalSurv showing the log-transformed mosquitoes per trap night for adult female *Culex tarsalis*. Values represent the mean number of mosquitoes per trap night for each county-year, for years from 2003 ('03') to 2023 ('23'). Raw data are shown in points, with nonlinear splines created using the 'loess' argument from geom_smooth() function.

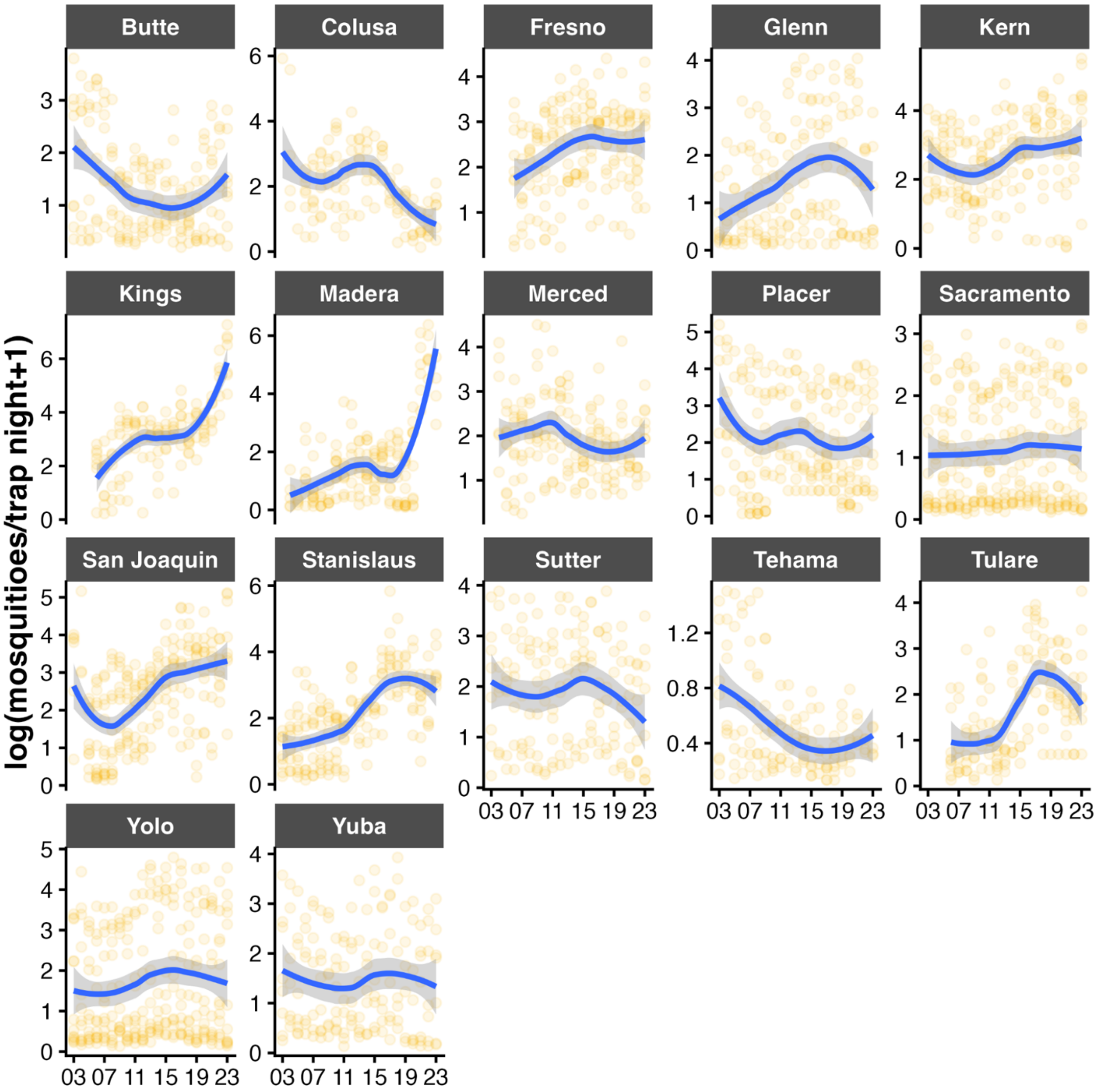

**Figure S5 Mosquito infection rate by county-year.** Data are from CalSurv showing log-transformed West Nile virus minimum infection rate (WNV MIR) per trap night for adult female *Culex tarsalis*, averaged by county and for years from 2003 (‘03’) to 2023 (‘23’). Raw data are shown in points, with nonlinear splines created using the ‘loess’ argument from geom_smooth() function.

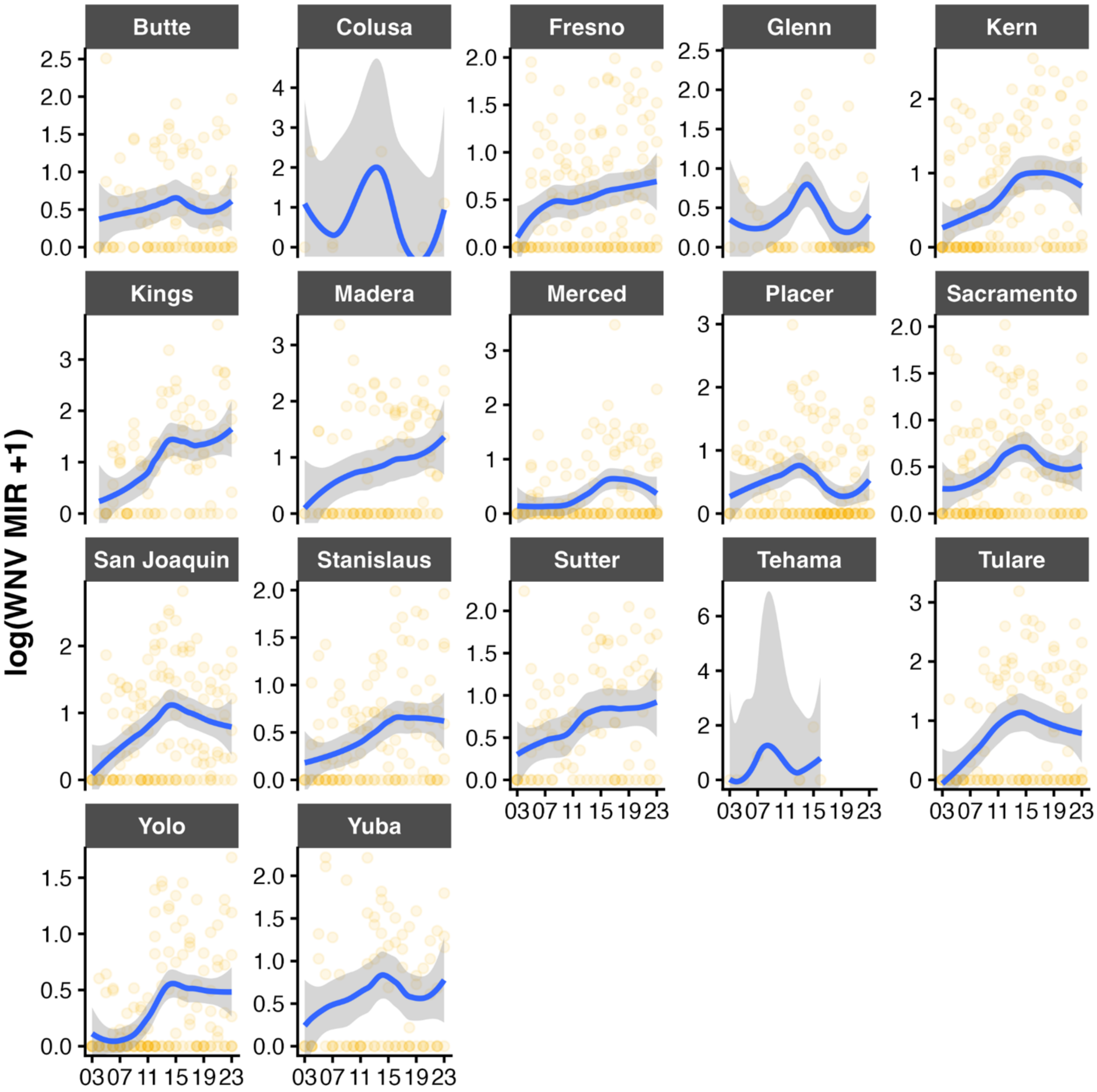

**Figure S6 Time series of hydrological variables.** Averaged Monthly values from 2003-2023. Variables shown are (a) soil moisture ($m^3/m^3$; CCI), (b) evapotranspiration (mm; GLEAM4), and standing water (% of water-covered pixels within 5 km buffer; GSW JRC) within 5 km of cluster unit. Nonlinear splines created using the 'loess' argument from geom_smooth() function.

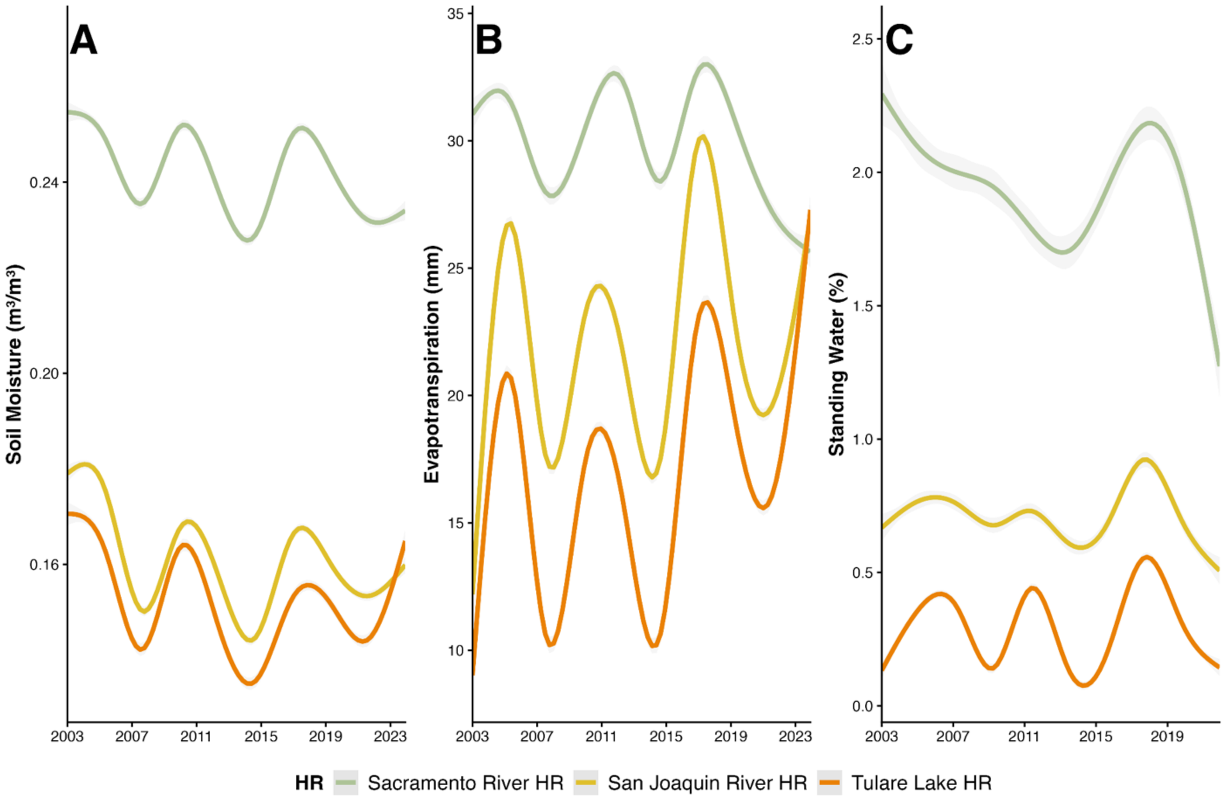

**Figure S7 Bird community competence across counties by year and month.** Averaged Monthly values from 2010-2023, when data was available for our study. Values represent the log-transformed bird community competence index, points are colored by county. Higher values represent higher bird community competence. Nonlinear temporal trends were modeled using a generalized additive model (GAM) from geom_smooth() function (k = 30).

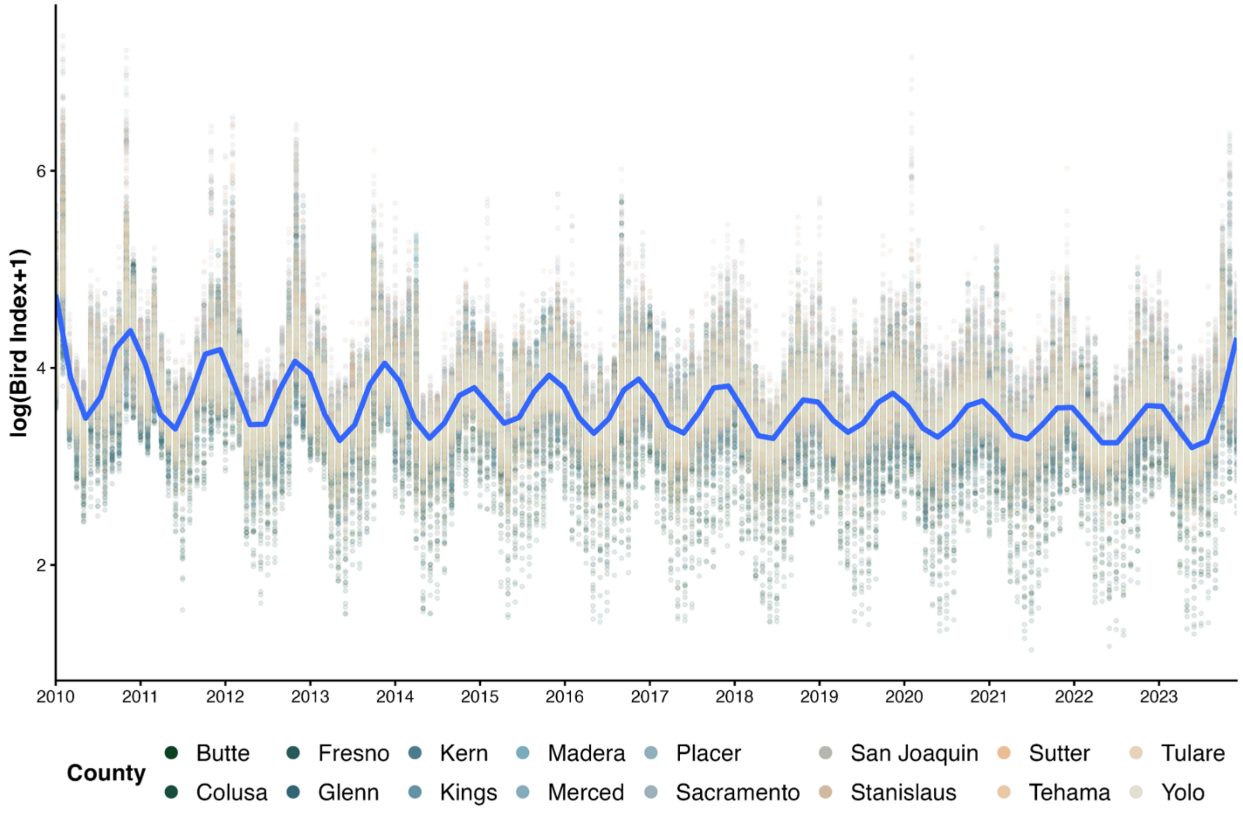

### Supplementary Tables

**Table S1 Data sources and resolutions.** All datasets used in this analysis, including their sources as well as their temporal and spatial resolutions, are reported. Temporal coverage is provided in the 'Notes' column where applicable. Additional details, including data citations, versions, and access links, are included in the footnotes.

| Variable | Source | Temporal resolution | Spatial resolution | Notes |
|---|---|---|---|---|
| **Mosquito abundance** | [1]CalSurv | Weekly | Trap stations | 2003-2023 |
| **Mosquito infection** | [1]CalSurv | Weekly | Trap stations | 2010-2023 |
| **Near-surface air temperature** | [2]CHELSA | Monthly; Daily | ~1 km | 2003-2021; 2022-2023 |
| **Palmer Drought Severity Index** | [3]gridMET | Monthly | ~4 km | 2003-2023 |
| **Actual Evapotranspiration** | [4]GLEAM | Monthly | ~10 km | |
| **Near-surface soil moisture** | [5]CCI | Monthly | ~25 km | Depth ~ 2 cm |
| **Standing water occurrence** | [6]JRC | Monthly | 30 m | |
| **Bird community competence index** | [7]eBird | Weekly | Coordinates | |
| **Precipitation** | [8]PRISM | 30-year normals | ~800 m | |
| **Irrigation method** | [9]Land IQ | Water year | Field-level | 2022 |
| **Spatial boundary of Central Valley** | [10]CDWR | 2012 | Polygon | |
| **Spatial boundary of hydro. regions** | [11]CDWR | 2019 | Polygon | |
| **Spatial boundary of major rivers** | [12]NHD | 2019 | Polygon | |
| **Spatial boundary of wetlands** | [13]NWI | 2004 | Polygon | |

Sources:

1. **California Vector-borne Disease Surveillance System** (CalSurv Gateway). *Mosquito surveillance data*. California Department of Public Health. Data request can be submitted to: https://ca.vectorsurv.org/.
2. **Climatologies at High Resolution for the Earth's Land Surface** (CHELSA) v2.1. *Temperature data*. Associated paper: (Karger et al. 2017) https://doi.org/10.1038/sdata.2017.122; Data available at: https://www.chelsa-climate.org/models/chelsa.
3. **Abatzoglou, J. T.** *gridMET: Gridded surface meteorological data.* Associated paper: (Abatzoglou 2013) *https://doi.org/10.1002/joc.3413;* Data available at: https://www.climatologylab.org/gridmet.html.
4. **Miralles, D.** et al. *GLEAM v4.2b evapotranspiration data.* Associated paper: (Miralles et al. 2016) https://doi.org/10.1038/s41597-025-04610-y; Data available at*: https://www.gleam.eu/*
5. **European Space Agency Climate Change Initiative.** *Soil moisture dataset.* Associated paper: https://ieeexplore.ieee.org/document/9165186/; Data available at: https://climate.esa.int/en/projects/soil-moisture/
6. **Joint Research Centre (JRC) Surface Water.** *Global Surface Water dataset.* Associated paper: (Preimesberger et al. 2021) https://doi.org/10.1038/nature20584; Data available at: https://global-surface-water.appspot.com/download.
7. **Cornell Lab of Ornithology.** *eBird Status and Trends data products.* Data available at: https://science.ebird.org/en/status-and-trends/data-access. *Index produced by MacDonald et al. 2024* https://doi.org/10.1002/ecy.4420.
8. **PRISM Climate Group, Oregon State University.** *PRISM climate data (normals).* Associated paper: (Abatzoglou 2013) https://doi.org/10.1002/joc.3413; Data available at: https://prism.oregonstate.edu/normals/.
9. **Land IQ.** *Land use and irrigation mapping data.* Data available at: https://www.landiq.com/land-use-mapping.
10. **California Department of Water Resources** (CDWR). Faunt, C.C. 2012. *Alluvial Boundary of California's Central Valley Survey data.* US Geological Survey. Data available at: https://www.sciencebase.gov/catalog/item/6314056bd34e36012efa2bf4.
11. **California Department of Water Resources** (CDWR). *Hydrological Region boundaries.* Spatial Data Standard v3.1. 2019. Data available at: https://gis.data.ca.gov/maps/3ce65cc69e0048169396b3610d6256b0/explore?location=36.824706%2C-119.742694%2C6
12. **National Hydrography Dataset** (NHD). US Geological Survey. Data available at: https://www.usgs.gov/national-hydrography/national-hydrography-dataset.
13. **National Wetlands Inventory** (NWI). US Fish and Wildlife Service. 2004. Data available at: https://www.fws.gov/program/national-wetlands-inventory.

**Table S2 Previous empirical *Culex*-climate studies.** Empirical studies (n > 48) examining associations between meteorological and hydrological variability and West Nile virus transmission or ecology in North America, with a focus on *Culex* spp. mosquitoes. Studies were included if they analyzed environmental variables derived from remote sensing products, gridded datasets, or weather/climate datasets (weather station-based studies were included when relevant). Only empirical analyses were retained; reviews and simulation studies were excluded. Vegetation and land cover variables (e.g., NDVI, EVI, NLCD classifications) were intentionally excluded to focus on atmospheric and hydrological drivers. Temporal aggregation choices (e.g., lagged variables, seasonable averages) are not distinguished in this table. Variables are grouped into major variable classes (e.g., precipitation, temperature) with similar variables (e.g., minimum/mean/maximum) consolidated for clarity. Study locations are indicated by state abbreviation or regional designations.

| Variable Class | Environmental Variable | Studies |
|---|---|---|
| **Drought and Hydrologic Anomaly Indices** | PDSI | Paull 2017 CO; Sambado 2025 CA; McMillan 2025 CT |
| | SPEI | Smith 2020 NE |
| | SPI | Smith 2020 NE |
| | PDI | Rosile & Bisesi 2017 OH |
| | PHDI | Skaff 2016 Midwestern US |
| **Precipitation** | Total precipitation | Ukawuba 2018 TX; Poh 2019 TX; Gizaw 2025 Western Canada; Davis 2018 SD; Smith 2020 NE; Trawinski & MacKay 2008 NY; Ward 2023 CA; Rosile & Bisesi 2017 OH; Wimberly 2014 Central US; Morin & Comrie 2013 Southern US; Keyel 2019 NY & CT; Soverow 2009 US; Shaman 2011 NY; Young 2013 US; Hess 2018 SD; Cooke 2006 MS; Kovach & Kilpatrick 2018 CA; Kovach & Kilpatrick 2024 CA; Winters 2008 CO; Paull 2017 CO; Ruiz 2010 IL; Gorris 2023 US; Johnson & Sukhdeo 2013 NJ; DeGroote 2008 IA; McMillan 2025 CT; Rhodes 2023 US; McMahon 2022 OK; Talbot 2024 Ontario Canada; Wimberly 2022 SD; Chuang 2012 SD; Karki 2016 IL; Hahn 2015 US; Chuang 2011 SD |
| | Precipitation anomalies | Keyel 2019 NY & CT; Soverow 2009 US; Hahn 2015 US |
| **Temperature** | Air temperature (minimum, mean, maximum) | Ukawuba 2018 TX; Poh 2019 TX; Gizaw 2025 Western Canada; Davis 2018 SD; Smith 2020 NE; Chuang 2012 Northern Great Plains; Kala 2017 CA; Trawinski & MacKay 2008 NY; Ward 2023 CA; Rosile & Bisesi 2017 OH; Liu & Weng 2012 CA; Wimberly 2014 Central US; Morin & Comrie 2013 Southern US; Keyel 2019 NY & CT; Soverow 2009 US; Shaman 2011 NY; Young 2013 US; Hess 2018 SD; MacDonald 2025 CA; Kovach & Kilpatrick 2018 CA; Kovach & Kilpatrick 2024 CA; Winters 2008 CO; Paull 2017 CO; Ruiz 2010 IL; Gorris 2023 US; Johnson & Sukhdeo 2013 NJ; Boser 2021 CA; DeGroote 2008 Iowa; McMillan 2025 CT; Rhodes 2023 US; McMahon 2022 OK; Talbot 2024 Ontario Canada; Martz 2025 AZ; Wimberly 2022 SD; Chuang 2012 SD; Gorris 2021 North America; Barker 2010 CA [*on Western Equine Encephalitis*]; Karki 2016 IL; Hahn 2015 US; Chuang 2011 SD |
| | Temperature anomalies | Keyel 2019 NY & CT; Hahn 2015 US |
| | Degree days (growing, cumulative, cooling) | Trawinski & MacKay 2008 NY; Keyel 2019 NY& CT; Winters 2008 CO; Paull 2017 CO [*number of weeks below freezing*]; Barker 2010 CA |
| **Atmospheric Moisture and Evaporative Demand** | Relative humidity | Gizaw 2025 Western Canada; Davis 2018 SD; Trawinski & MacKay 2008 NY; Hess 2018 SD; MacDonald 2025 CA; Winters 2008 CO; McMahon 2022 OK; Wimberly 2022 SD; Chuang 2012 SD; Karki 2016 IL; Chuang 2011 SD |
| | Specific humidity | Ukawuba 2018 TX; Ward 2023 CA; Shaman 2011 NY; Gorris 2021 North America |
| | VPD | Davis 2018 SD; Hess 2018 SD; Wimberly 2022 SD |

| | | |
|---|---|---|
| | Evapotranspiration (actual, potential, total) | Davis 2018 SD; Chuang 2012 Northern Great Plains; Kala 2017 CA; Trawinski & MacKay 2008 NY; Ward 2023 CA; Morin & Comrie 2013 Southern US; Cooke 2006 MS |
| | Dew point temperature | Soverow 2009 US; DeGroote 2008 IA |
| **Surface Energy Balance** | Surface pressure | Ward 2023 CA |
| | Surface skin temperature | Ward 2023 CA |
| **Surface Hydrology and Water Availability** | Soil moisture **Ponding frequency from Soil Survey Geographic Database soils data* | Ukawuba 2018 TX; Davis 2018 SD; Chuang 2012 SD*; Trawinski & MacKay 2008 NY*; Hess 2018 SD *; Ward 2023 CA; Keyel 2019 NY & CT; Shaman 2011 NY; Chuang 2012 SD |
| | Surface water presence or extent, including wetlands and streams (metrics include presence, extent, density, distance) | Chuang 2012 SD; Kala 2017 CA; Trawinski & MacKay 2008 NY; Morin & Comrie 2013 Southern US; Hess 2018 SD; Cooke 2006 MS; Skaff 2016 Midwestern US; MacDonald 2025 CA; DeGroote 2008 IA; Liu 2008 IN; Chuang 2012 SD |
| | River discharge **River gauges* | Sambado 2025 CA; Reisen 1992 CA*; Wegbreit & Reisen 2000* |
| | Irrigation (presence, distance) | Chuang 2012 SD [*water draw point distance*]; Kovach & Kilpatrick 2018 CA; Kovach & Kilpatrick 2024 CA; Cardenas 2011 TX [*flooding canals*]; DeGroote 2008 IA; Gates 2009 US [*census-level irrigation*]; L Eisen 2010 CO |
| | Normalized difference water index (NDWI) | Liu & Weng 2012 CA |
| | Hydrological model | Shaman & Day 2005 FL; Cooke 2006 MS; Shaman 2005 FL |
| | Normalized difference moisture index (NDMI) | McMahon 2022 OK |
| **Cryosphere** | Snow (snow cover, SWE, snowmelt timing) | Winters 2008 CO |
| **Large-scale Climate Modes** | El Niño-Southern Oscillation (ENSO) | Poh 2019 TX |
| **Atmospheric Dynamics** | Wind Speed | Gizaw 2025 Western Canada; Karki 2016 IL |

*Sources:* (Reisen et al. 1992, Wegbreit and Reisen 2000, Shaman et al. 2005, 2011, Shaman and Day 2005, Cooke et al. 2006, DeGroote et al. 2008, Liu et al. 2008, Winters et al. 2008, Gates and Boston 2009, Soverow et al. 2009, Ruiz et al. 2010, Barker et al. 2010, Trawinski and Mackay 2010, BARKER et al. 2010, Chuang et al. 2011, 2012b, 2012a, Cardenas et al. 2011, Liu and Weng 2012, Chuang and Wimberly 2012, Johnson and Sukhdeo 2013, Morin and Comrie 2013, Young et al. 2013, Wimberly et al. 2014, 2022, 2022, Hahn et al. 2015, Karki et al. 2016, Skaff and Cheruvelil 2016, Rosile and Bisesi 2017, Paull et al. 2017, Kala et al. 2017, Ukawuba and Shaman 2018, Kovach and Kilpatrick 2018, 2024, Davis et al. 2018, Hess et al. 2018, Keyel et al. 2019, 2019, Poh et al. 2019, Smith et al. 2020, Gorris et al. 2021, 2023, Boser et al. 2021, McMahon et al. 2022, Rhodes et al. 2023, Ward et al. 2023, Talbot et al. 2024, Gizaw et al. 2025, MacDonald et al. 2025, McMillan et al. 2025, Sambado et al. 2025, Martz et al. 2025)

**Table S3 Variation in mosquitoes and hydrological variables.** Extended summary statistics for *Culex tarsalis* surveillance data (April-October, 2003-2023). Counties are grouped by hydrological region: Sacramento River (SAC), San Joaquin River (SAN), and Tulare Lake (TUL). Sampling effort is summarized as the total number of clusters and mosquitoes (trap nights). County-level vector and environmental conditions are reported as mean (standard deviation): West Nile virus minimum infection rate (WNV MIR), Palmer Drought Severity Index (PDSI), near-surface Soil Moisture (SM, $m^{3/}m^3$), standing surface water (SW, % of pixels with water occurrence), and actual Evapotranspiration (ET, mm).

| County | Clusters | Mosquitoes (trap nights) | WNV MIR | PDSI | SM | SW (5 km) | ET |
|---|---|---|---|---|---|---|---|
| **Butte** (SAC) | 43 | 651,748 (14,497) | 2.67 (5.8) | -1.22 (2.9) | 0.18 (0.07) | 3.19 (7.10) | 40.29 (30.86) |
| **Colusa** (SAC) | 7 | 106,465 (2,128) | 2.24 (6.0) | -1.30 (3.0) | 0.18 (0.07) | 5.74 (8.90) | 32.30 (28.96) |
| **Fresno** (SAN) | 412 | 1,096,600 (14,050) | 3.10 (9.7) | -0.99 (3.0) | 0.12 (0.05) | 0.60 (1.09) | 19.76 (23.01) |
| **Glenn** (SAC) | 13 | 361,359 (5,125) | 2.76 (4.8) | -1.36 (3.1) | 0.17 (0.07) | 4.61 (8.43) | 34.91 (29.99) |
| **Kern** (TUL) | 181 | 2,395,117 (21,395) | 3.54 (10.3) | -0.71 (2.9) | 0.11 (0.04) | 0.52 (1.39) | 10.50 (14.47) |
| **Kings** (TUL) | 70 | 699,184 (5,374) | 5.84 (14.0) | -0.93 (2.9) | 0.11 (0.06) | 0.33 (0.60) | 16.67 (21.12) |
| **Madera** (SAN) | 108 | 452,135 (13,844) | 6.46 (12.4) | -0.96 (3.1) | 0.11 (0.05) | 0.30 (0.70) | 22.48 (26.36) |
| **Merced** (SAN) | 109 | 514,133 (24,410) | 2.99 (16.8) | -1.07 (3.0) | 0.09 (0.07) | 0.80 (1.98) | 21.78 (24.91) |
| **Placer** (SAC) | 130 | 596,310 (9,252) | 1.90 (13.6) | -0.81 (2.7) | 0.18 (0.07) | 3.49 (8.98) | 36.90 (30.06) |
| **Sacramento** (SAC) | 264 | 1,074,754 (88,418) | 4.82 (33.1) | -1.06 (2.9) | 0.24 (0.08) | 2.34 (4.53) | 30.19 (26.38) |
| **San Joaquin** (SAN) | 124 | 976,250 (12,797) | 4.20 (22.4) | -1.28 (2.9) | 0.14 (0.05) | 2.31 (4.67) | 30.99 (27.98) |
| **Stanislaus** (SAN) | 176 | 712,704 (12,733) | 2.22 (23.2) | -1.07 (2.8) | 0.12 (0.05) | 0.63 (1.25) | 26.25 (27.07) |
| **Sutter** (SAC) | 33 | 891,833 (11,951) | 3.61 (8.65) | -1.27 (3.0) | 0.16 (0.07) | 3.46 (5.43) | 30.24 (27.91) |
| **Tehama** (SAC) | 13 | 21,650 (2,103) | 2.13 (4.5) | -1.15 (2.8) | 0.16 (0.06) | 0.87 (0.71) | 36.72 (30.93) |
| **Tulare** (TUL) | 198 | 330,804 (15,449) | 4.43 (13.6) | -0.97 (3.1) | 0.10 (0.05) | 0.23 (0.46) | 19.79 (22.05) |
| **Yolo** (SAC) | 102 | 2,561,061 (64,146) | 1.78 (8.9) | -1.18 (3.1) | 0.21 (0.06) | 2.36 (4.10) | 26.45 (26.51) |
| **Yuba** (SAC) | 17 | 316,028 (10,355) | 2.86 (5.22) | -1.20 (2.9) | 0.16 (0.07) | 3.59 (4.48) | 36.57 (29.06) |

### Supplementary Text

#### Text S1 Additional details on mosquito surveillance and data processing.

*Mosquito surveillance data*. Data for this project were obtained from the CalSurv Gateway through data request (1) #000092, approved on 15 January 2026, and (2) #000098 approved on 17 March 2026 by the California Vectorborne Disease Surveillance System. Surveillance covered the following California counties: Butte, Colusa, Fresno, Glenn, Kern, Kings, Madera, Merced, Placer, Sacramento, San Joaquin, Shasta, Sutter, Tehama, Tulare, Yolo, and Yuba. Data was requested for the years 2003-2023. Due to the bird community competence index being available for only 2010-2023, we only look at WNV MIR data for the same years to include bird community competence in our infection models but keep abundance models for 2003-2023.

*Mosquito cleaning and filtering*. Abundance data were restricted to female mosquitoes (bloodfed, gravid, mixed, or unfed), with recorded species names, classified as an abundance collection type and without trap malfunctions. Data were further subset to the months of April through October, representing peak mosquito activity. Only *Culex tarsalis*, *Culex pipiens*, *Culex quinquefasciatus* were retained. After filtering, the dataset included 38,592,919 individual female *Culex* mosquitoes collected over 2,832,101 trap nights (1,232,452 recorded observations) . A total of 36,770 unique trap stations were aggregated into 2,902 spatial clusters. Virus testing data were also restricted to female mosquitoes of the three primary *Culex* species. Testing targets (West Nile virus) were kept if test status was reported confirmed or negative. After filtering, the dataset included 557,840 unique testing pools (1,038,287 recorded observations) from 18,491 unique trap stations. Our analysis and visualizations in the main text focused only on *Cx. tarsalis*, but we kept *Cx. pipiens* and *Cx. quinquefasciatus* observations for comparisons in Supplementary Text 8.

*Mosquito outcomes.* Mosquito outcomes, abundance and infection rates, were calculated per mosquito species-trap-year-month and then averaged per spatial cluster. Abundance data was calculated as the average number of adult female mosquitoes per trap night. Pools of mosquito samples were screened for West Nile virus infection using the equation: MIR = ((number of positive pools / number of mosquitoes tested) * 1000). A small fraction of observations exhibits very high MIR values (> 100), which arise mechanistically when positive pools are detected among very small numbers of tested mosquitoes. Because these values reflect instability in the denominator rather than meaningful variation in infection intensity, we conduct robustness checks excluding MIR > 100 observation (approximately 0.13% of the sample). Results were qualitatively unchanged (Supplementary Text 8).

**Text S2 Additional details on hydrological variables for variable rationale and data processing.**

*(i) Palmer Drought Severity Index (PDSI)*
Rationale: The PDSI is a widely used indicator of long-term landscape "wetness" and drought severity. It captures the balance of precipitation and evapotranspiration over extended periods and is relevant for mosquito ecology because drought conditions can reduce breeding habitats and alter mosquito-host interactions. Previous studies have shown that increased drought severity can reduce mosquito abundance but increase West Nile virus transmission, potentially due to enhanced mosquito-host aggregation around limited bodies of water (Shaman et al. 2005, 2011, Sambado et al. 2025). We hypothesize that the effects of PDSI on mosquitoes and WNV may vary spatially, depending on county-level hydrological characteristics such as flood regimes, proximity to water bodies (e.g., wetlands, major rivers), and water management practices.

Data Processing: Monthly PDSI data were obtained from the gridMET gridded climate dataset at a ~ 4 km resolution (Abatzoglou 2013) with `climateR` package (cite: mikejohnson51/climateR). Values were extracted for each spatial cluster and averaged to monthly means per cluster. Positive values indicate wetter-than-average conditions, while negative values indicate drier-than-average conditions. Typical values range from -4 (extreme drought) to 4 (extremely wet conditions).

*(ii) Soil Moisture*
Rationale: Surface soil moisture (top few centimeters of soil) represents localized water availability and potential mosquito breeding habitats, such as ephemeral pools. Increased soil moisture can expand breeding opportunities and potentially boost adult mosquito abundance. However, excessive moisture may flush eggs or reduce larval survival. Additionally, we expect that soil moisture effects on mosquito populations may exhibit temporal lags, reflecting developmental times from egg to adult.

Data processing: We used a harmonized, bias-corrected soil moisture dataset from the European Space Agency Climate Change Initiative (CCI) program v09.2 (Dorigo et al. 2017, Gruber et al. 2019, Preimesberger et al. 2021). CCI v09.2 is a global dataset spanning 1978 to 2023 and is based on satellite data. The COMBINED product integrates active and passive microwave satellite observations to provide near-surface soil moisture (~0-5 cm) at a 0.25° (~ 25 km) spatial resolution. Daily observations were aggregated to monthly means, and cluster-level averages were calculated. Data are reported in volumetric units ($m^3/m^3$) and accessed in NetCDF-4 format on 21 February 2026.

*(iii) Standing Surface Water*
Rationale: Standing surface water reflects local ecological conditions, influencing both mosquito populations and host communities. Low water availability may concentrate mosquitoes and bird hosts, potentially increasing contact rates and WNV transmission. Conversely, abundant water can either enhance mosquito breeding by providing more habitat or reduce populations if large water bodies support predators of mosquito larvae.

Data processing: We used the Global Surface Water Dataset (European Commission Joint Research Centre (GSW JRC)) derived from Landsat imagery (Pekel et al. 2016). GSW JRC is a product of Landsat satellite images available from 1984-2021. Standing surface water occurrence is defined as the monthly frequency of water presence per 30 m pixel. To capture local- and landscape-scale variation, we calculated the % of water-covered pixels within a 5 km buffer around each cluster centroid. For each monthly image, water presence was converted to a binary mask, where pixels classified as seasonal or permanent (class 2 or 3) were coded as water (1), and all other pixels were classified as 0. We then calculated the proportion of water-covered pixels within the 5 km buffer and then multiplied by 100 to express water coverage as a percentage of the total buffer area. As sensitivity analysis, we also did this for 10 km buffers which we keep in the supplemental files but focus our main analysis on 5 km buffer. Data was processed in Google Earth Engine and were accessed on 22 February 2026.

*(iv) Evapotranspiration.*
Rationale: Total actual evapotranspiration (ET) integrates soil evaporation, plant transpiration, and canopy interception loss, and serves as a proxy for atmospheric and vegetation moisture. Higher ET may enhance adult mosquito survival by reducing desiccation stress and supporting microhabitats (Boser et al. 2021). We hypothesize that increased ET promotes higher adult mosquito populations.

Data processing: ET estimates (Et, mm $day^{-1}$) were obtained from the Global Land Evaporation Amsterdam Model (GLEAM v4.2b) (Miralles et al. 2025). GLEAM4.2b is a global dataset spanning 2003 to 2024 and is based on satellite data and is considered an observation-driven model. We used the monthly aggregated Et variable, which includes soil, plant, and canopy contributions at 0.1° (~10 km) spatial resolution. Cluster-level averages were calculated, with values reported in mm per month. Data were accessed in NetCDF-4 format on 21 February 2026.

**Text S3 Other data sources and justifications.**

*(i) Precipitation.* Precipitation (ppt; mm) was derived from gridded climate data provided by the Parameter-elevation Regressions on Independent Slopes Model (PRISM Group, Oregon State University, https://prism.oregonstate.edu, accessed 16 March 2026) (Abatzoglou 2013, Daly et al. 2015). Thirty-year precipitation normals (1991-2020) at an 800 m spatial resolution were used to visualize the spatial patterns in annual total precipitation (Fig. 1a) and seasonal variation in mean monthly precipitation (Fig. 1b).

*(ii) Wetland extent.* Wetland data were obtained from the US Geological Survey National Wetland Inventory (https://www.fws.gov/program/national-wetlands-inventory) and subsetted for California. Spatial boundaries were processed to quantify the wetland distribution within the study area. Polygon features were first extracted for the California Central Valley and clipped to the study extent. To reduce file size and improve computational efficiency, the resulting dataset was exported to GeoPackage format and geometrically simplified using a 100 m tolerance while preserving polygon topology. Wetland features with small spatial extent (shape area < 5,000 $m^2$) were excluded to reduce noise associated with highly fragmented features. The filtered dataset was further simplified using rmapshaper (ms_simplify; keep = 0.02, preserve shapes = TRUE) to retain overall spatial structure while reducing vertex density. The final dataset was used to visualize wetland presence and extend within California's Central Valley (Fig. 1c), representing habitat availability for *Culex* mosquito populations.

*(iii) Irrigation type classification.* Cropland spatial data was acquired from Land IQ ( https://www.landiq.com/land-use-mapping), developed in collaboration with the California Department of Water Resources (CADWR), which provides field-level crop classification across California. For this study, crop types were reclassified into three primary irrigation categories: flood, sprinkler, and drip irrigation. Crop group codes (identified by prefixes F, I, P, G, T, YP, X, D, C, V, R) were mapped to irrigation types based on their predominant irrigation practices. Specifically, crop groups were classified as follows: flood (R, P), sprinkler (G, F), drip (V, C, D, T, YP), and unknown (I, X) (Fig. 1d). These assignments were based on the irrigation method most commonly associated with each crop group. Although some crops may be irrigated using multiple methods, we assigned a single dominant irrigation type (e.g., flooded conditions for rice) for consistency; future work could explore mixed irrigation practices in greater detail. These three irrigation categories were selected based on their different implications for mosquito habitat availability. Flood irrigation generates extensive standing water bodies that are strongly associated with *Culex tarsalis* oviposition habitats. Sprinkler irrigation typically results in more transient surface water due to evaporation and runoff, potentially creating short-lived breeding opportunities. Drip irrigation is the most water-efficient method and generally limits standing water, although small, localized pools may persist near the base of crops (e.g., orchards), which could still support mosquito development.

*(iv) Temperature.* Air temperature data were obtained from the CHELSA v2.1 global climate dataset (https://www.chelsa-climate.org/models/chelsa, accessed 16 March 2026) (Karger et al. 2017). Monthly mean near-surface air temperature (tas) rasters were downloaded for 2003-2021. Data were accessed programmatically from the CHELSA monthly archive. For 2022-2023, when

monthly products were not available, daily CHELSA temperatures were obtained using the `Rchelsa` package (https://github.com/inSilecoBlogLegacy/rchelsa) and downloaded in batches for the study extent. Daily rasters were aggregated to monthly means to match the monthly dataset. All temperature values were converted from Kelvin to degrees Celsius and spatially averaged within the sampling clusters using zonal extraction using the `terra` package (Hijmans 2024). The resulting data provides a continuous monthly time series of air temperature across 2003-2023 used for panel models.

*(v) Bird community competence.* Birds are key amplifying hosts in the West Nile virus (WNV) transmission cycle, but species differ in their ability to acquire, maintain, and transmit infection to *Culex* mosquitoes. In addition, bird community composition varies seasonally due to migration, leading to spatial and temporal variation in host competence. To account for these processes, we used a previously developed bird community competence index (MacDonald et al. 2024) which integrates species-specific reservoir competence with spatiotemporally explicit estimates of bird community composition. Briefly, species abundance was modeled at a ~ 1 km resolution using eBird observational data (Fink et al. 2025) for the top 20 WNV-competent bird species (Kilpatrick et al. 2007). These species-level abundance surfaces were combined with published estimates to generate a monthly, raster-based community competence index. The resulting index was then spatially aggregated to study clusters by averaging raster values within each polygon, producing a monthly estimate of host community competence used in panel models.

**Text S4. Main mosquito abundance panel model**

**S4.1 Specification and identification**
We estimate the effect of Palmer Drought Severity Index (PDSI) on mosquito abundance using panel regression models with ordinary least squares fixed effects, implemented via the `fixest` package in R (Bergè 2019). The analysis uses monthly data from 2003-2023 (N = 21 years) across 1,863 spatial clusters (N = 82,078 observations). All models include cluster, year, and month fixed effects, and standard errors are computed using Conley corrections with a 5 km spatial cutoff to account for spatial dependency across nearby clusters.

The outcome is log-transformed mosquito abundance per trap night to reduce right-skewness and allow coefficients to be interpreted approximately as percent changes. The primary predictor PDSI, a standardized index of landscape moisture conditions typically ranges from -4 (dry) to 4 (wet). Control variables include mean air surface temperature, modeled as both a linear and quadratic term to capture nonlinear biological responses of the mosquito and virus. All covariates are standardized. Month fixed effects are defined over the seven-month peak mosquito surveillance season (April-October).

The baseline specification is:

$$Y_{iym} = \beta PDSI_{iym} + \beta X_{iym} + \alpha_i + \gamma_y + \delta_m + \varepsilon_{iym}$$

where *i* indexes spatial clusters, *y* year, and *m* month. $Y_{iym}$ is log-transformed mosquito abundance, $X_{iym}$ is the vector of control variables, *i* are cluster fixed effects, *y* are year fixed effects, and *m* are month fixed effects. $\varepsilon_{iym}$ the error term.

Identification relied on within-cluster temporal variation in PDSI after removing time-invariant spatial differences and common interannual and seasonal shocks. The remaining deviations in PDSI within a cluster over time are the estimates of interest. We assume that PDSI reflects broad scale hydroclimatic conditions that are unaffected by local mosquito dynamics at monthly time scales.

**S4.2 Main results**
Our baseline model indicates that wetter conditions (higher PDSI values) are significantly associated with higher mosquito abundance ($\beta$ = 0.15, standard error (SE) = 0.02, 95% confidence intervals (CI) [0.12, 0.19], $p < 0.001$). This corresponds to an approximately 16.5% increase in mosquito abundance per one-unit increase in PDSI, controlling for temperature and fixed effects. A one-unit change in PDSI reflects a meaningful shift in overall landscape moisture conditions rather than short-term weather variability. Temperature showed a nonlinear relationship with mosquito abundance. Marginal effects are positive at lower temperatures, peaks at approximately 29°C, and declines at higher temperatures, consistent with biologically constrained mosquito development and virus survival. The model explains a moderate share of variation after accounting for fixed effects, with RMSE = 0.95, overall adjusted $R^2$ = 0.53, and within $R^2$ = 0.008. The relatively low $R^2$ reflects substantial residual short-term variation in mosquito abundance at the cluster-month level after absorbing spatial and temporal fixed effects, which is typical in ecological panel settings.

**S4.3 Diagnostics**

**S4.3.1 Fixed-effect robustness.** We assess the sensitivity of estimated effects to alternative fixed-effects structures, ranging from pooled OLS (no fixed effects; FE) to progressively more restrictive specifications, including cluster fixed effects, cluster-by-month fixed effects, and cluster-by-year fixed effects. This table outlines the estimated associations between PDSI and mosquito abundance under different dimensions of unobserved heterogeneity (spatial, seasonal, or interannual variation).

| Variables | Pooled OLS | Cluster FE | Cluster, Month FE | Cluster, Year FE | Cluster, Year, Month FE |
|---|---|---|---|---|---|
| **PDSI** | 0.12***<br>(0.009) | 0.12***<br>(0.008) | 0.15***<br>(0.018) | 0.15***<br>(0.017) | 0.15***<br>(0.017) |
| **Temp.** | 1.02***<br>(0.055) | 0.80***<br>(0.042) | 0.76***<br>(0.039) | 0.17***<br>(0.047) | 0.17***<br>(0.047) |
| **Temp.$^2$** | -0.34***<br>(0.031) | -0.20***<br>(0.020) | -0.18***<br>(0.021) | -0.22***<br>(0.022) | -0.22***<br>(0.022) |
| **Obs.** | 82,212 | 82,078 | 82,078 | 82,078 | 82,078 |
| **$R^2$** | 0.058 | 0.52 | 0.53 | 0.54 | 0.54 |
| **Within $R^2$** | | 0.091 | 0.078 | 0.008 | 0.008 |

***p < 0.001. FE, fixed effects. Temp., temperature. Conley standard errors (5 km).

**S4.3.2 Multicollinearity.** We assess multicollinearity among covariates using variance inflation factors (VIFs), calculated using the \`car\` package (Fox and Weisberg 2019). As expected, PDSI exhibits minimal collinearity with temperature controls, suggesting that hydroclimatic variation is distinct from local thermal conditions. The linear and quadratic temperature terms exhibit moderate VIF values due to their functional relationship. Temperature controls are standardized (std).

| PDSI | Temperature (std) | Temperature (std)$^2$ |
|---|---|---|
| 1.01 | 7.20 | 7.18 |

**S4.4 Robustness checks.** We conduct a set of robustness checks to evaluate the sensitivity of the baseline estimate for mosquito abundance. These analyses assess whether the estimated association between PDSI and abundance is robust to alternative assumptions regarding: (i) unobserved heterogeneity, (ii) functional form and distributional assumptions, (iii) temporal persistence in hydroclimatic effects, (iv) temperature specification, and (v) spatial dependence in inference.

All models are estimated using Conley standard errors with a 5 km spatial cutoff unless otherwise specified (see S4.4.5). Reported coefficients represent the effect of PDSI on abundance, where positive values indicate wetter landscape conditions and negative values indicate drier conditions. Confidence intervals are reported at the 95% level.

**S4.4.1 Alternative fixed effects.** We first assess robustness to alternative specifications of unobserved heterogeneity by varying the fixed-effect structure. This tests whether the estimated PDSI effect is sensitive to different assumptions about spatial, seasonal, and interannual confounding.

We estimate models including: (i) cluster-by-month-by-year fixed effects, (ii) cluster-by-year fixed effects, (iii) cluster-by-month fixed effects, (iv) cluster fixed effects, and (v) pooled OLS without fixed effects. This sequence progressively relaxes controls for time-invariant and time-varying unobserved heterogeneity. Consistency of the PDSI coefficient across specifications supports the interpretation that the baseline estimate is driven by within-cluster temporal variation rather than omitted variables.

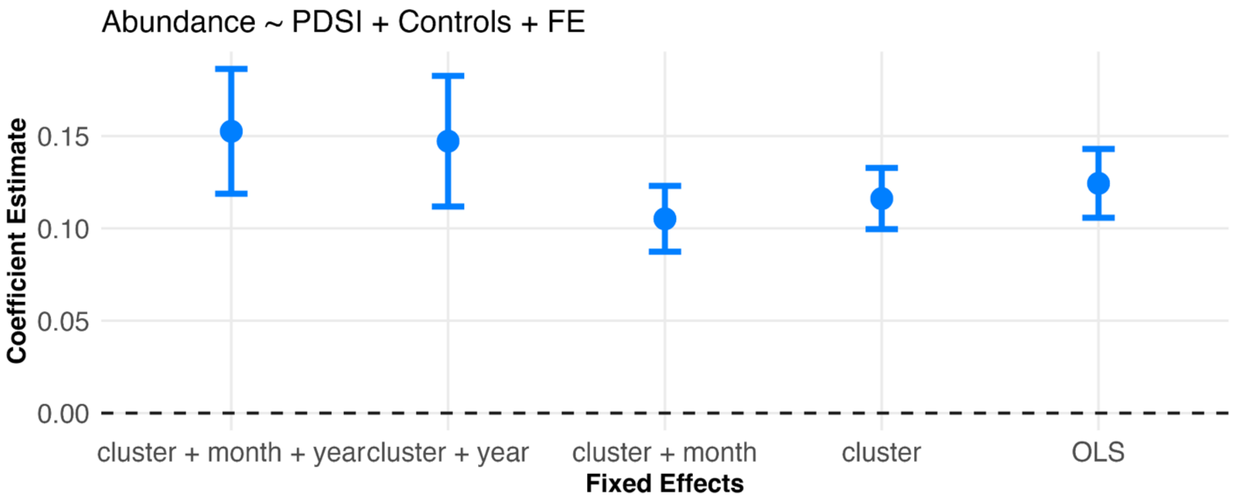


**S4.4.2 Model type.** We evaluate robustness to alternative distributional assumptions by estimating the baseline model using different regression frameworks appropriate for count and semi-continuous outcomes. Specifically, we compare log-linear models (baseline specification using log-transformed abundance), Poisson models, and negative binomial models using the feols(), fepois(), and fenegbin() functions, respectively (Bergè 2019). Stability across specifications indicates that inference is not driven by transformation choices or distributional misspecification.

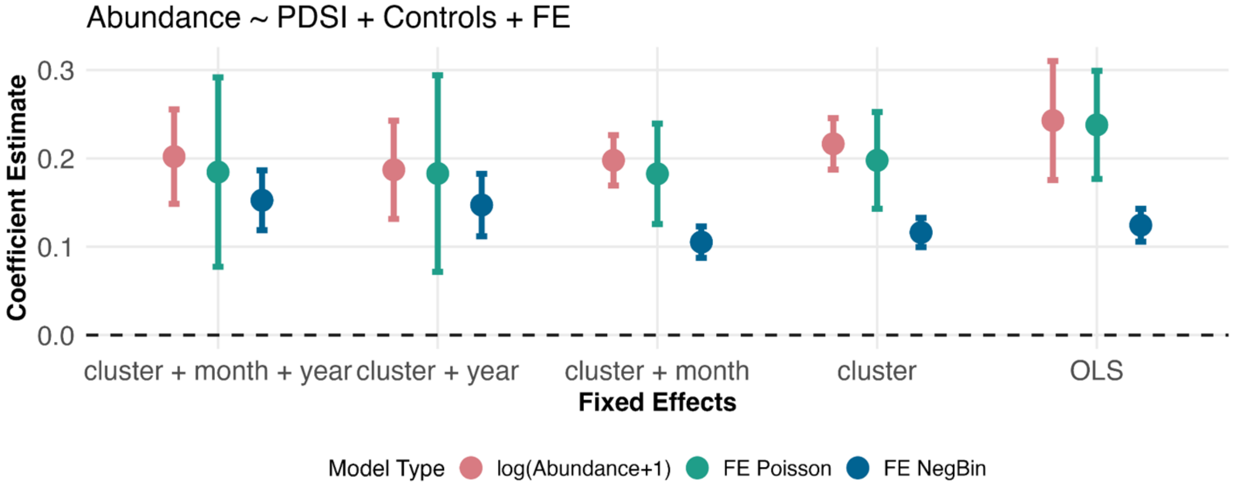

**S4.4.3 PDSI lags.** To assess temporal dynamics in landscape wetness response, we estimate models using lagged values of PDSI ranging from contemporaneous conditions (0-month lag) to 5-month lag structures. The moderate robust effects across lag structures indicate that the PDSI-mosquito relationship is not an artifact of a particular time lag, which is expected since PDSI is supposed to represent long-term averages rather than short responses.

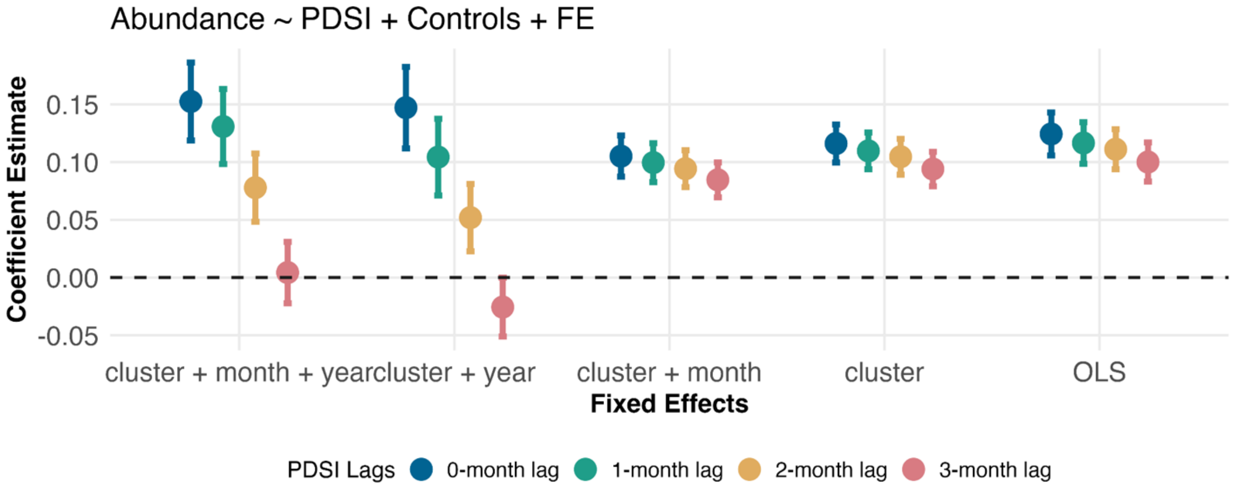


**S4.4.4. Temperature specification.** We examine sensitivity to alternative functional forms of temperature controls, which account for nonlinearities in mosquito development and virus transmission. Models include (i) no temperature controls, (ii) linear temperature term, and (iii) a quadratic term specification. Stability of PDSI coefficients across these specifications suggest that estimated effects of PDSI are not impacted by temperature functional form assumptions.

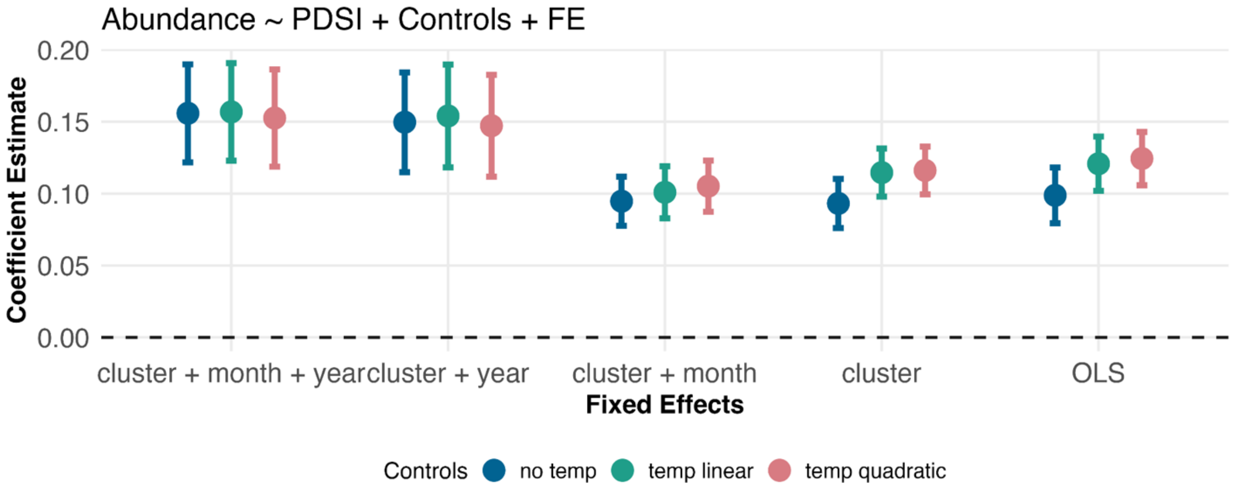


**S4.4.5 Conley spatial cutoffs.** We evaluate robustness of inference to alternative assumptions about spatial correlation in the error structure. We re-estimate models using Conley standard errors with spatial cutoffs ranging from 1.5 km to 25 km (1.5, 5, 10, 15, 20, and 25 km). Across all cutoffs, coefficient estimates remain significant. We ultimately chose 5 km cutoffs as a reasonable value for mosquito dispersal pattern in the California's Central Valley.

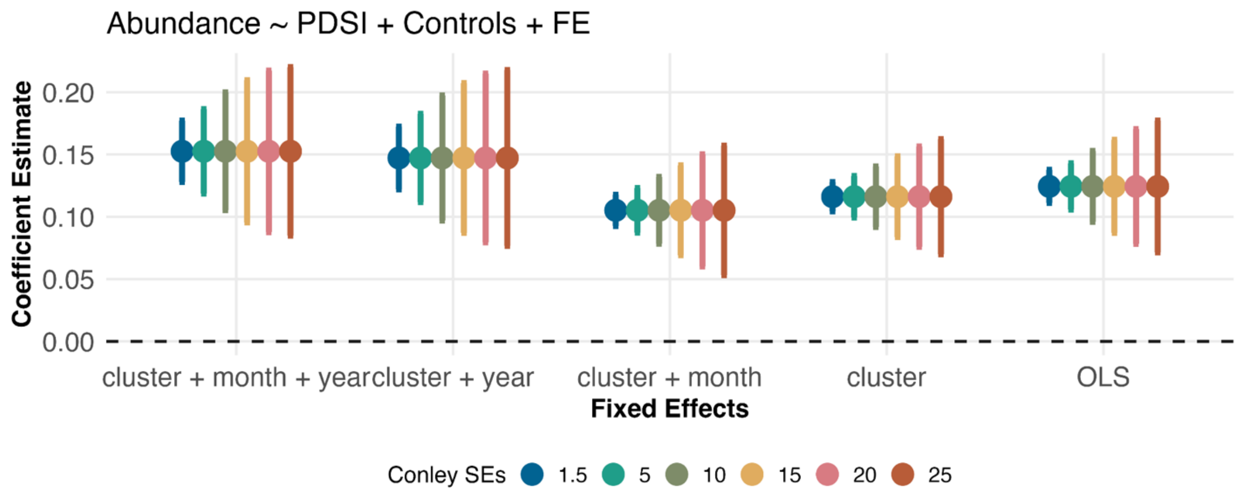


**S4.5 Supporting figures.** We present supporting figures to visualize key distributional properties and functional form assumptions underlying the main specification. First, we show histograms of raw mosquito abundance per trap night along its log-transformed distribution. These figures illustrate the strong right-skewness in the raw outcomes and the resulting normalization achieved through the log transformation. All model estimates are therefore reported using log(mosquitoes per trap night + 1) to reduce skewness and improve interpretability of coefficients as approximate percentage changes.

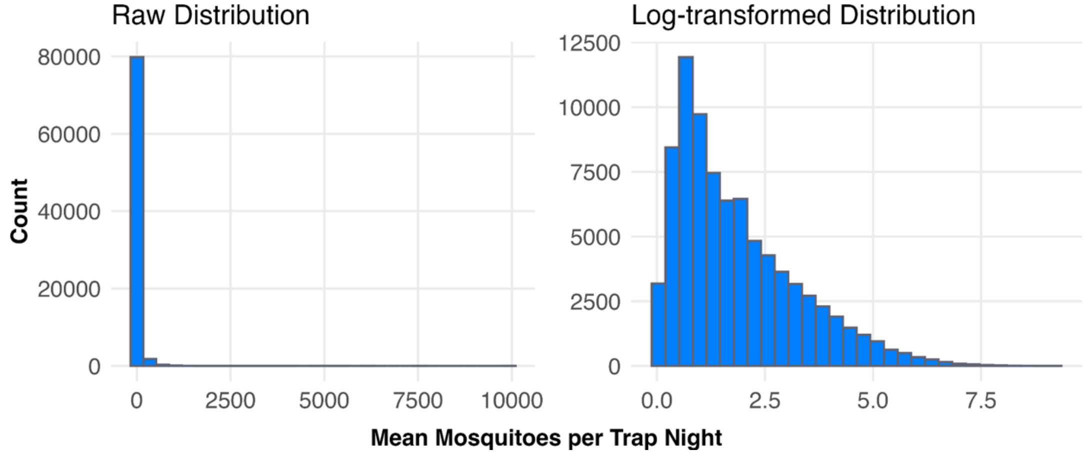


Next, we show marginal effect plots for temperature to assess the assume function form. This figure highlights the nonlinearity in the relationship between mosquito abundance and temperature.

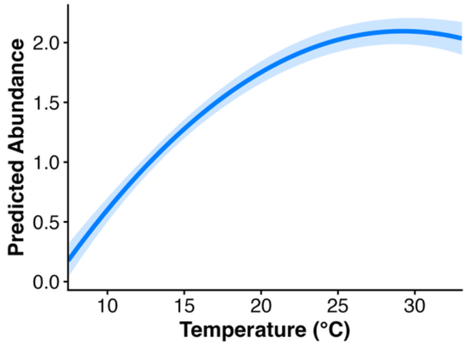

### Text S5. Main mosquito infection panel model

#### S5.1 Specification and identification

We estimate the effect of Palmer Drought Severity Index (PDSI) on West Nile virus minimum infection rate (WNV MIR) using panel regression models with ordinary least squares fixed effects, implemented via the `fixest` package in R (Bergè 2019). The analysis uses monthly data from 2010-2023 (N = 14 years) across 1,272 spatial clusters (N = 25,302 observations). All models include cluster, year, and month fixed effects, and standard errors are computed using Conley corrections with a 5 km spatial cutoff to account for spatial dependency across nearby clusters.

The outcome is log-transformed WNV MIR to reduce right-skewness and allow coefficients to be interpreted approximately as percent changes. Due to extreme outliers, we cut off values of MIR WNV  100, which resulted in 0.13% data loss. We demonstrate the robustness of this cut off choice in S5.6. The primary predictor PDSI, a standardized index of landscape moisture conditions typically ranges from -4 (dry) to 4 (wet). Control variables include mean air surface temperature, modeled as both a linear and quadratic term to capture nonlinear biological responses of the mosquito and virus. We also include a bird community competence index to represent the relative abundance of potentially infectious blood meal sources. All covariates are standardized. Month fixed effects are defined over the seven-month peak mosquito surveillance season (April-October).

The baseline specification is:

$$Y_{iym} = \beta PDSI_{iym} + \beta X_{iym} + \alpha_i + \gamma_y + \delta_m + \varepsilon_{iym}$$

where *i* indexes spatial clusters, *y* year, and *m* month. $Y_{iym}$ is log-transformed WNV MIR, $X_{iym}$ is the vector of control variables, *i* are cluster fixed effects, *y* are year fixed effects, and *m* are month fixed effects. $\varepsilon_{iym}$ the error term.

Identification relied on within-cluster temporal variation in PDSI after removing time-invariant spatial differences and common interannual and seasonal shocks. The remaining deviations in PDSI within a cluster over time are the estimates of interest. We assume that PDSI reflects broad scale hydroclimatic conditions that are unaffected by local mosquito virus dynamics at monthly time scales.

#### S5.2 Main results

Our baseline model indicates that wetter conditions (higher PDSI values) are significantly associated with lower mosquito infection rates (WNV MIR) ($\beta$ = -0.053, SE = 0.014, 95% CI [-0.08, -0.03], p < 0.001). This corresponds to an approximately 5.4% decrease in mosquito infection rate per one-unit increase in PDSI, controlling for temperature, bird community structure, and fixed effects. A one-unit change in PDSI reflects a meaningful shift in overall landscape moisture conditions rather than short-term weather variability. Temperature showed a nonlinear relationship with mosquito infection rates. Marginal effects are positive at lower temperatures, peaks at approximately 29 °C, and declines at higher temperatures, consistent with biologically constrained mosquito development and virus survival. Marginal effects of bird community competence showed a linear relationship where an increase in bird community competence led to an increase in WNV MIR. The model explains a moderate share of variation after accounting for fixed effects, with RMSE = 0.82, overall adjusted $R^2$ = 0.17, and within $R^2$ = 0.0030. The low $R^2$ reflects substantial residual short-term variation in mosquito WNV MIR at the cluster-month level after absorbing spatial and temporal fixed effects, which is typical in ecological panel settings. We also expected infection models to be more biologically complex than abundance models, hence more difficult to explain higher variation.

#### S5.3 Diagnostics

**S5.3.1 Fixed-effect robustness.** We assess the sensitivity of estimated effects to alternative fixed-effects structures, ranging from pooled OLS (no fixed effects) to progressively more restrictive specifications, including cluster fixed effects, cluster-by-month fixed effects, and cluster-by-year fixed effects. This table outlines the estimated associations between PDSI and mosquito WNV MIR under different dimensions of unobserved heterogeneity (spatial, seasonal, or interannual variation).

| Variables | Pooled OLS | Cluster FE | Cluster, Month FE | Cluster, Year FE | Cluster, Year, Month FE |
|---|---|---|---|---|---|
| **PDSI** | 0.01*<br>(0.005) | 0.005<br>(0.006) | -0.03*<br>(0.014) | -0.05***<br>(0.014) | -0.05***<br>(0.014) |
| **Temp.** | 0.23***<br>(0.045) | 0.28***<br>(0.042) | 0.21***<br>(0.037) | -0.088<br>(0.058) | -0.088<br>(0.058) |
| **Temp.$^2$** | 0.15***<br>(0.032) | 0.11***<br>(0.028) | 0.18***<br>(0.024) | 0.15***<br>(0.033) | 0.145***<br>(0.033) |
| **Bird Index** | 0.09***<br>(0.013) | 0.086***<br>(0.013) | 0.093***<br>(0.014) | -0.004<br>(0.016) | -0.004<br>(0.016) |
| **Obs.** | 25,518 | 25,302 | 25,302 | 25,302 | 25,302 |
| **$R^2$** | 0.055 | 0.15 | 0.16 | 0.21 | 0.21 |
| **Within $R^2$** | | 0.051 | 0.059 | 0.003 | 0.003 |

***p < 0.001; *p<0.05. FE, fixed effects. Temp., temperature. Bird Index, bird community competence index. Conley standard errors (5 km).

**S5.3.2 Multicollinearity.** We assess multicollinearity among covariates using variance inflation factors (VIFs), calculated using the `car` package (Fox and Weisberg 2019). As expected, PDSI and bird community competence index (bird index) exhibits minimal collinearity with temperature controls, suggesting that hydroclimatic variation is distinct from local thermal conditions. The linear and quadratic temperature terms exhibit moderate VIF values due to their functional relationship. Temperature and bird controls are standardized (std).

| PDSI | Temperature (std) | Temperature (std)$^2$ | Bird Index |
|---|---|---|---|
| 1.01 | 9.96 | 9.97 | 1.05 |

**S5.4 Robustness checks.** We conduct a set of robustness checks to evaluate the sensitivity of the baseline estimate for mosquito WNV MIR. These analyses assess whether the estimated association between PDSI and WNV MIR is robust to alternative assumptions regarding: (i) unobserved heterogeneity, (ii) functional form and distributional assumptions, (iii) temporal persistence in hydroclimatic effects, (iv) temperature specification, and (v) spatial dependence in inference.

All models are estimated using Conley standard errors with a 5 km spatial cutoff unless otherwise specified (see S5.4.5). Reported coefficients represent the effect of PDSI on WNV MIR, where positive values indicate wetter landscape conditions and negative values indicate drier conditions. Confidence intervals are reported at the 95% level.

**S5.4.1 Alternative fixed effects.** We first assess robustness to alternative specifications of unobserved heterogeneity by varying the fixed-effect structure. This tests whether the estimated PDSI effect is sensitive to different assumptions about spatial, seasonal, and interannual confounding.

We estimate models including: (i) cluster-by-month-by-year fixed effects, (ii) cluster-by-year fixed effects, (iii) cluster-by-month fixed effects, (iv) cluster fixed effects, and (v) pooled OLS without fixed effects. This sequence progressively relaxes controls for time-invariant and time-varying unobserved heterogeneity. Consistency of the PDSI coefficient across specifications supports the interpretation that the baseline estimate is driven by within-cluster temporal variation rather than omitted variables. For infection models, fixed effects appear to be important in identifying the effect on WNV MIR compared to ordinary least squares where the relationship is slightly positive.

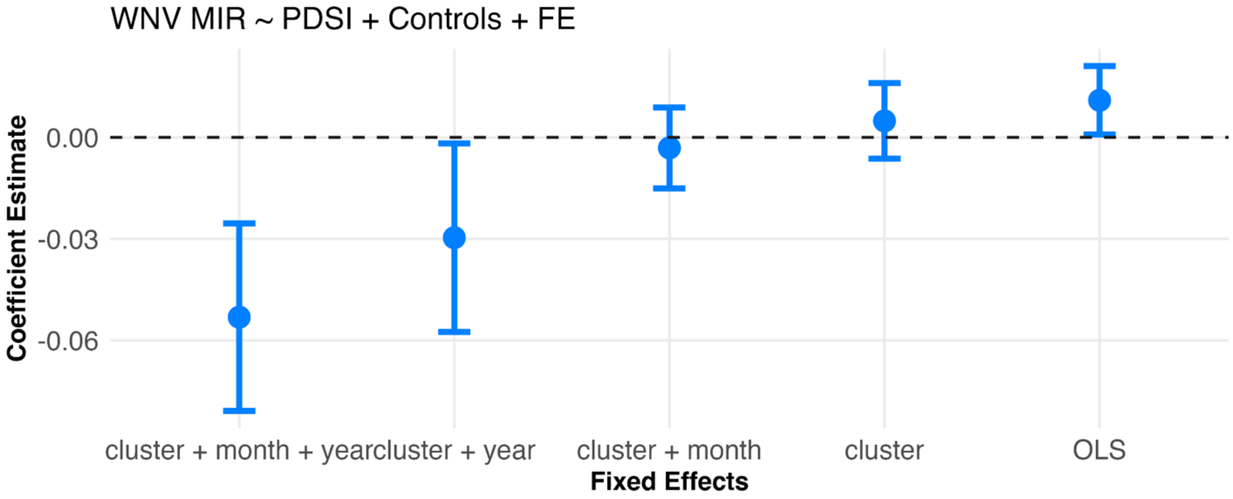


**S5.4.2 Model type.** We evaluate robustness to alternative distributional assumptions by estimating the baseline model using different regression frameworks appropriate for count and semi-continuous outcomes. Specifically, we compare log-linear models (baseline specification using log-transformed abundance), Poisson models, and negative binomial models using the feols(), fepois(), and fenegbin() functions, respectively (Bergè 2019). Stability across specifications indicates that inference is not driven by transformation choices or distributional misspecification.

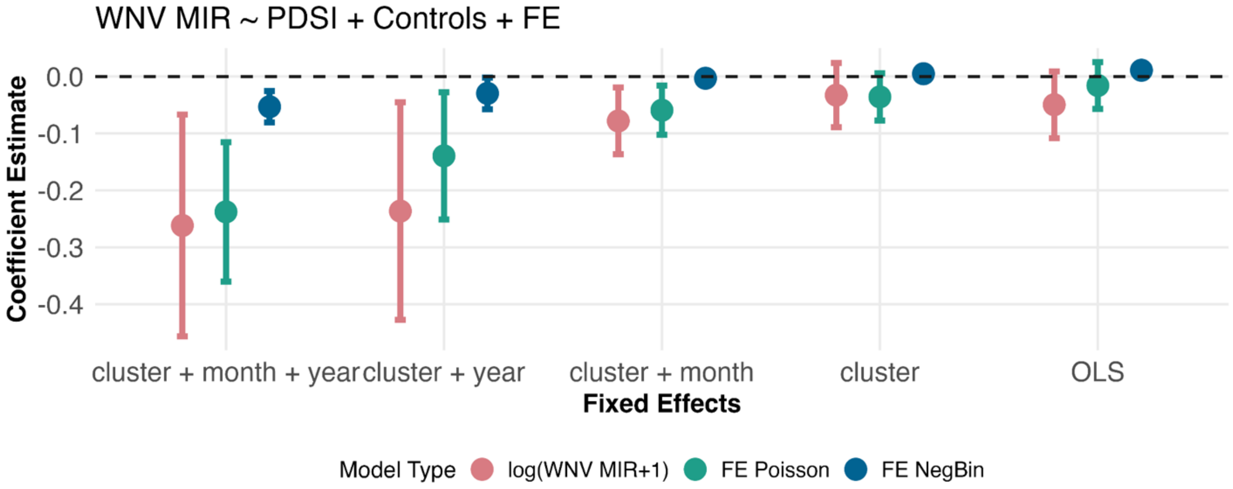


**S5.4.3 PDSI lags.** To assess temporal dynamics in landscape wetness response, we estimate models using lagged values of PDSI ranging from contemporaneous conditions (0-month lag) to 5-month lag structures. The moderate robust effects across lag structures indicate that the PDSI-mosquito relationship is not an artifact of a particular time lag, which is expected since PDSI is supposed to represent long-term averages rather than short responses.

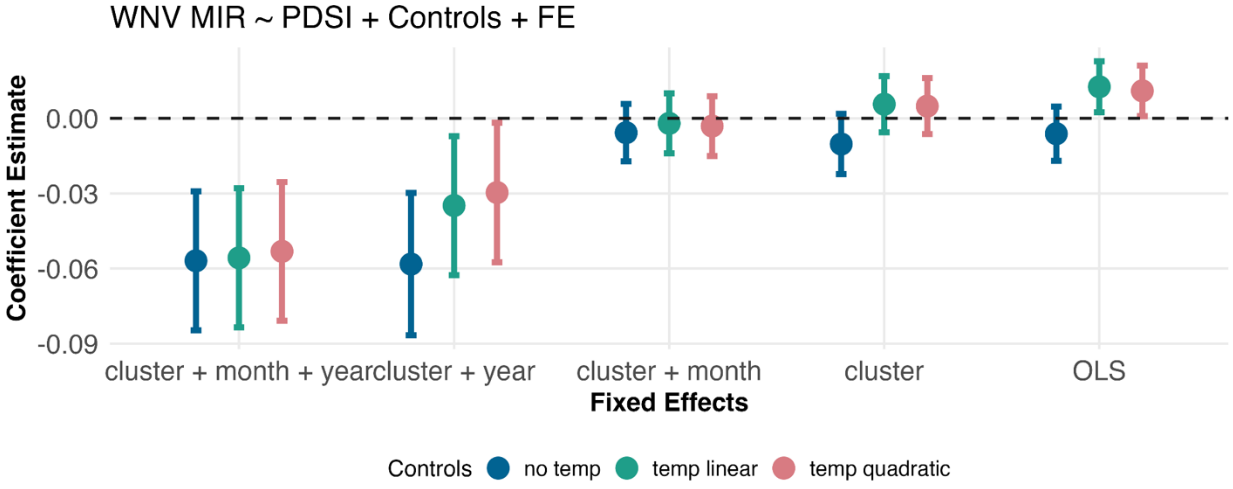


**S5.4.4. Temperature specification.** We examine sensitivity to alternative functional forms of temperature controls, which account for nonlinearities in mosquito development and virus transmission. Models include (i) no temperature controls, (ii) linear temperature term, and (iii) a quadratic term specification. Stability of PDSI coefficients across these specifications suggest that estimated effects of PDSI are not impacted by temperature functional form assumptions.

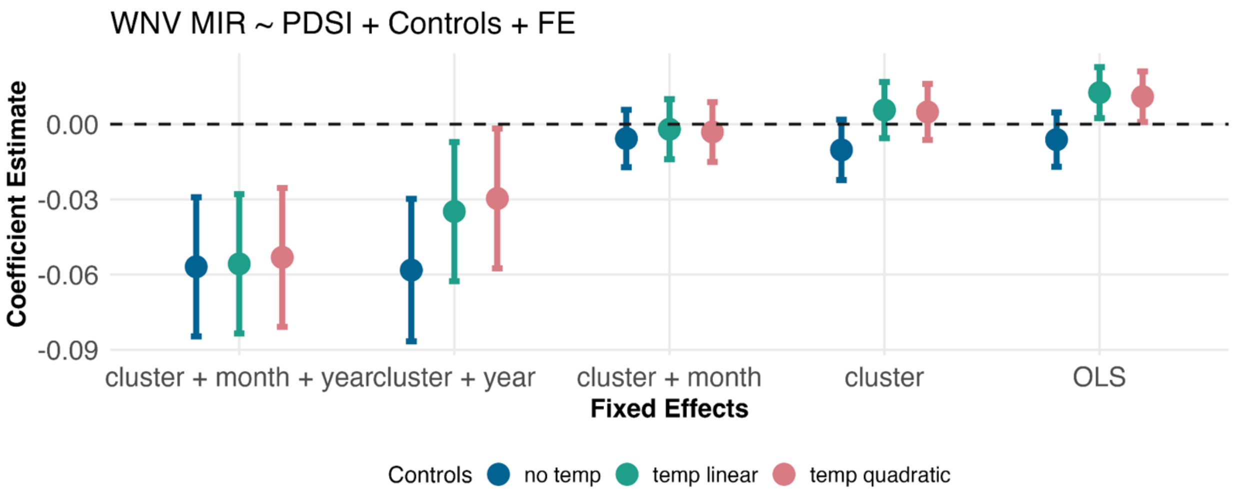


**S5.4.5 Conley spatial cutoffs.** We evaluate robustness of inference to alternative assumptions about spatial correlation in the error structure. We re-estimate models using Conley standard errors with spatial cutoffs ranging from 1.5 km to 25 km (1.5, 5, 10, 15, 20, and 25 km). Across all cutoffs, coefficient estimates remain significant. We ultimately chose 5 km cutoffs as a reasonable cutoff of mosquito dispersal pattern in our region.

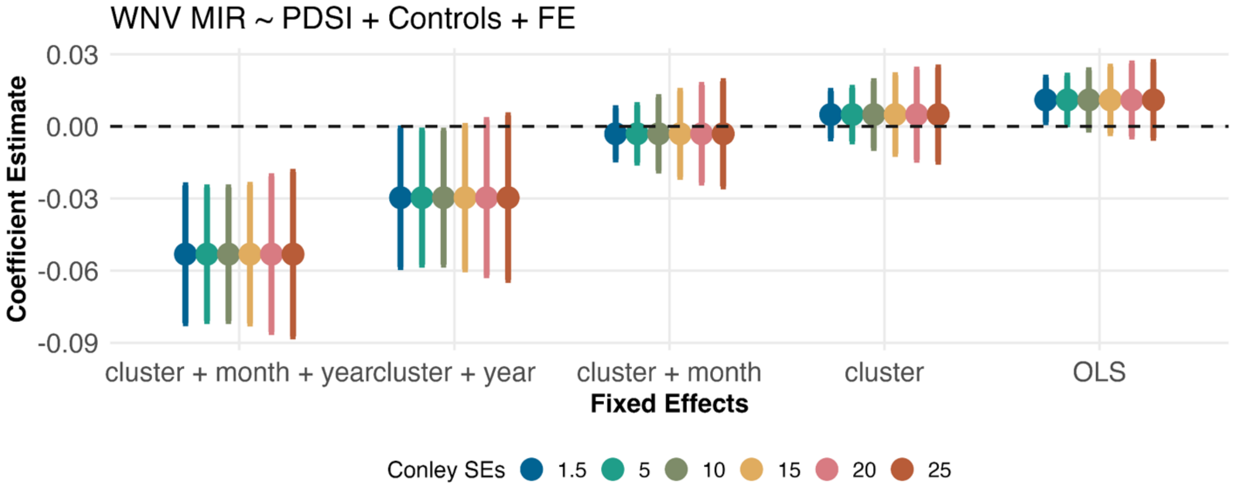


**S5.4.6 WNV MIR cutoffs.** We assess robustness to high-end values of WNV MIR by comparing (i) the full sample including all observations up to the observed maximum (MIR 1000), (ii) a sample excluding observations at the maximum observed value (MIR < 1000), (iii) a sample excluding high MIR values (MIR 100), and (iv) a specification in which MIR values are capped at 100. These alternative constructions evaluate sensitivity of the estimated associations to upper-tail WNV MIR values driven by small-denominator instability in the MIR measure.

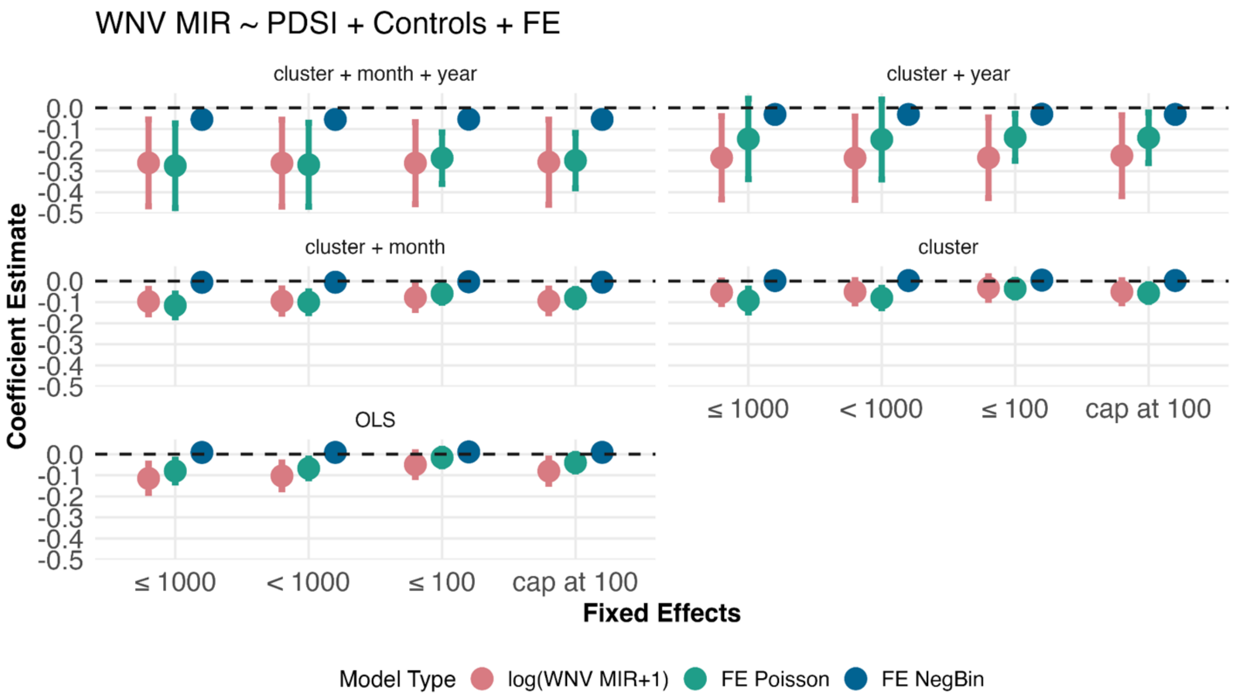


**S5.5 Supporting figures.** We present supporting figures to visualize key distributional properties and functional form assumptions underlying the main specification. First, we show histograms of raw mosquito WNV MIR along its log-transformed distribution. These figures illustrate the strong right-skewness in the raw outcomes and the resulting normalization achieved through the log transformation. All model estimates are therefore reported using log(WNV MIR + 1) to reduce skewness and improve interpretability of coefficients as approximate percentage changes.

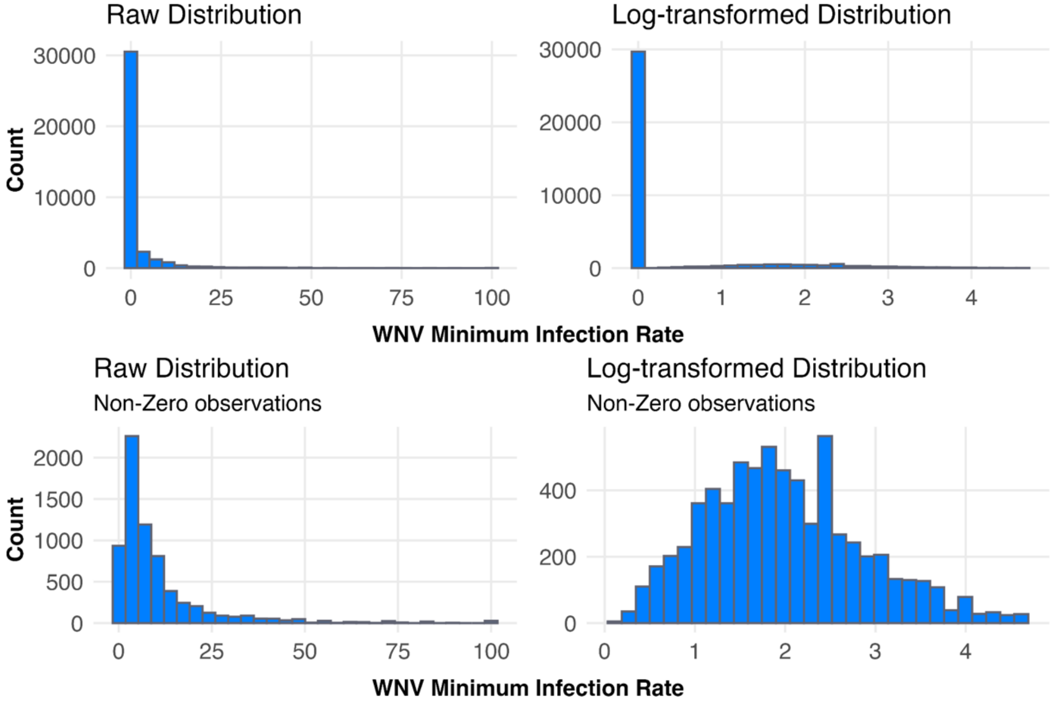

Next, we show marginal effect plots for temperature and bird community competence index. This figure highlights the nonlinearity in the relationship between mosquito WNV MIR and temperature. The relationship between WNV MIR and bird community competence is positively linear.

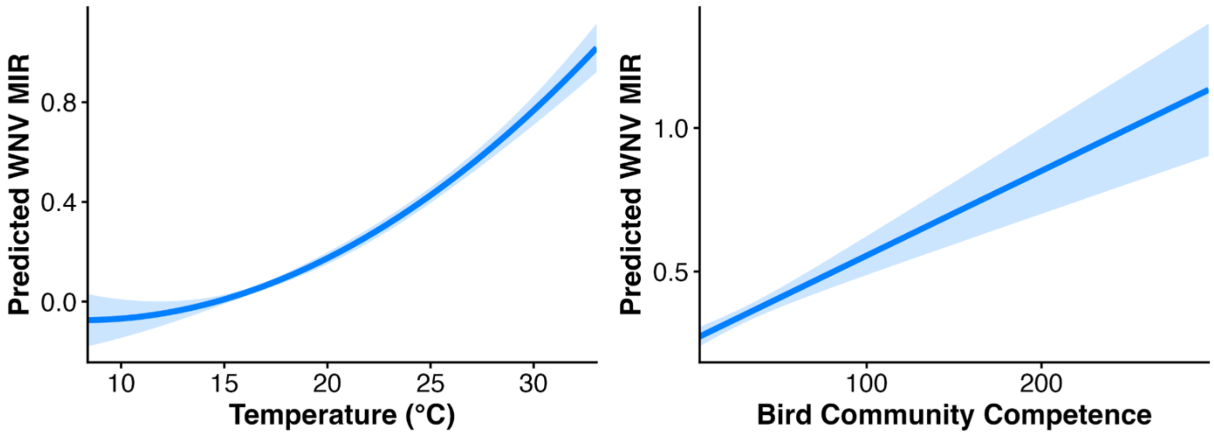

**Text S6. Spatial heterogeneity panel models**

**S6.1 Abundance models**

We extend the baseline abundance model (Text S4) to allow the effect of PDSI to vary across hydrological regions by interacting PDSI with region indicators. The model is estimated on the same panel dataset (2003-2023) using ordinary least squares with cluster, year, and month fixed effects. Standard errors are computed using Conley correction with a 5 km spatial cutoff. Control variables follow the baseline specification, including temperature and its quadratic form.

Hydrological regions are used as the primary spatial grouping because they reflect ecologically meaningful differences in surface hydrology and water management, which are likely to influence mosquito habitat availability and population dynamics. As a robustness check, we also estimated analogous models using county-level interactions (17 counties in the Central Valley) to assess finer-scale spatial variation in the PDSI-mosquito relationship.

The estimating equation is:

$$Y_{iym} = \beta(PDSI_{iym} * Region_r) + \beta X_{iym} + \alpha_i + \gamma_y + \delta_m + \varepsilon_{iym}$$

where *r* represents the region-specific marginal effect of PDSI.

**S6.1.1 Results**

Estimated marginal effects of PDSI on mosquito abundance vary across hydrological regions. All region-specific coefficients are positive and statistically significant ($p < 0.001$), indicating that wetter conditions are associated with higher mosquito abundance. Model fit is similar to the baseline model, with RMSE = 0.95, overall adjusted $R^2 = 0.53$, and within $R^2 = 0.014$.

| **Hydrological Regions** | **Estimate** | **95% CI** | **p-value** |
|---|---|---|---|
| **Sacramento River** | 0.06 | 0.024, 0.10 | **< 0.001** |
| **San Joaquin River** | 0.16 | 0.12, 0.20 | **< 0.001** |
| **Tulare Lake** | 0.25 | 0.20, 0.30 | **< 0.001** |

**Observations** = 82,078

County-specific marginal effects are computed as the sum of the baseline PDSI coefficient and the corresponding county interaction term, yielding the total effect of PDSI within each county. The figure presents the resulting county-level estimate with 95% confidence intervals. The results show spatial heterogeneity, although most counties show positive associations between wetter conditions and mosquito abundance. Counties within the same hydrological region tend to cluster in similar ranges, but within-region variation persists, highlighting heterogeneity at fine spatial scales.

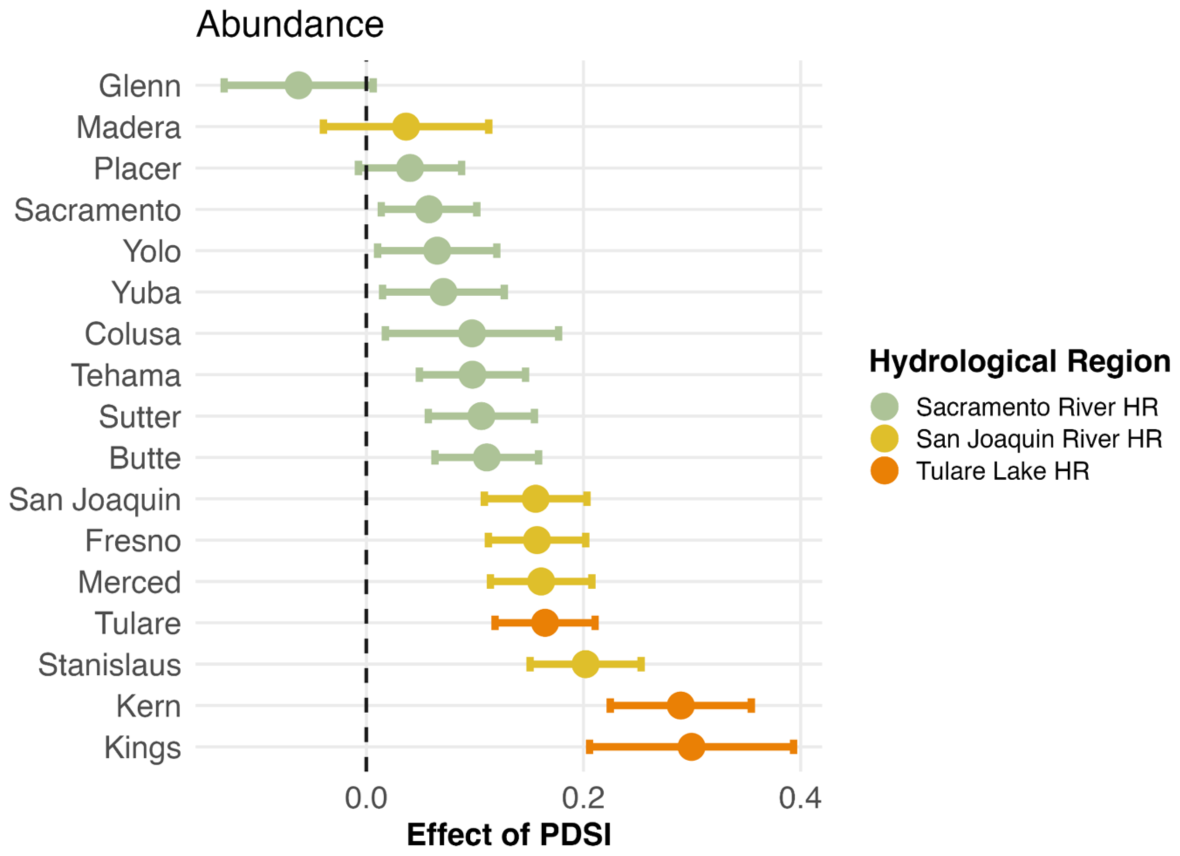


**S6.2 WNV MIR models**

We extend the baseline abundance model (Text S5) to allow the effect of PDSI to vary across hydrological regions by interacting PDSI with region indicators. The model is estimated on the same panel dataset (2003-2023) using ordinary least squares with cluster, year, and month fixed effects. Standard errors are computed using Conley correction with a 5 km spatial cutoff. Control variables follow the baseline specification, including temperature and its quadratic form, and bird community competence index.

Hydrological regions are used as the primary spatial grouping because they reflect ecologically meaningful differences in surface hydrology and water management, which are likely to influence mosquito habitat availability and population dynamics. As a robustness check, we also estimated analogous models using county-level interactions (17 counties in the Central Valley) to assess finer-scale spatial variation in the PDSI-mosquito relationship.

The estimating equation is:

$$Y_{iym} = \beta(PDSI_{iym} * Region_r) + \beta X_{iym} + \alpha_i + \gamma_y + \delta_m + \varepsilon_{iym}$$

where *r* represents the region-specific marginal effect of PDSI.

**S6.2.1 Results**

Estimated marginal effects of PDSI on mosquito WNV MIR vary across hydrological regions. All region-specific coefficients are positive and statistically significant ($p < 0.001$), indicating that wetter conditions are associated with lower mosquito infection rates. Model fit is similar to the baseline model, with RMSE = 0.84, overall adjusted $R^2$ = 0.16, and within $R^2$ = 0.003.

| **Hydrological Regions** | **Estimate** | **95% CI** | **p-value** |
|---|---|---|---|
| **Sacramento River** | -0.06 | -0.09, -0.03 | **< 0.001** |
| **San Joaquin River** | -0.04 | -0.07, -0.01 | **0.02** |
| **Tulare Lake** | -0.08 | -0.12, -0.04 | **< 0.001** |
| **Observations** = 25,340 | | | |

County-specific marginal effects are computed as the sum of the baseline PDSI coefficient and the corresponding county interaction term, yielding the total effect of PDSI within each county. The figure presents the resulting county-level estimate with 95% confidence intervals. The results show spatial heterogeneity, although most counties show negative associations between wetter conditions and mosquito infection rates. There is less spatial clustering of counties within the same hydrological region, including non-significant relationships.

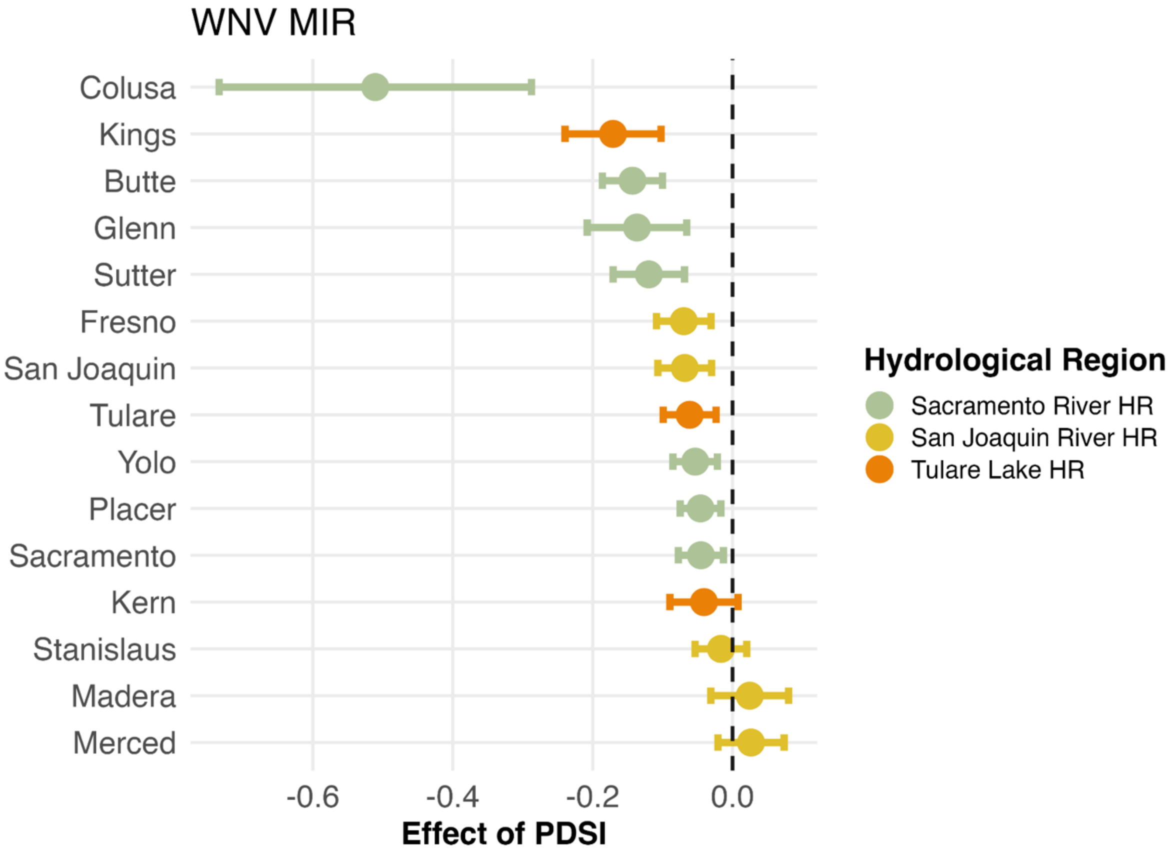

**Text S7. Hydrological panel models**

To identify which components of water availability are most strongly associated with mosquito abundance and WNV infection, we re-estimate the baseline fixed-effects model (Text S4, S5) replacing PDSI with alternative hydrological variables, including soil moisture, evapotranspiration, and standing surface water (% of water covered pixels within 5- and 10-km buffers). Each hydrological variable is estimated in separate models to avoid multicollinearity and is included with distributed lags from 0 to 3 months. All models include cluster, year, and month fixed effects and are estimated using OLS with Conley-corrected standard errors (5 km cutoff). Covariates follow the baseline specification, and all hydrological variables are standardized.

The model is:

$$Y_{iym} = \beta HydroVar_{iym,l} + \beta X_{iym} + \alpha_i + \gamma_y + \delta_m + \varepsilon_{iym}$$

where *l* denotes the month lag of 0, 1, 2, or 3. Month lag 0 refers to contemporaneous month.

#### S7.1 Abundance models results

Results are reported in the table below, which present coefficient estimates and standard errors (SE) for each hydrological variable and lag. Each row corresponds to a separate regression. Positive coefficients indicate that higher values of the hydrological variable are associated with higher mosquito abundance at the specified month lag. Because each hydrological variable is estimated in separate regressions, coefficients should be interpreted as independent associations conditional on fixed effects and covariates. All covariates were standardized.

| Hydrological Variable | Month Lag | Estimate | SE |
|---|---|---|---|
| **Soil Moisture** | 0 | -0.016 | 0.041 |
| **Evapotranspiration** | | 0.091*** | 0.014 |
| **Surface water (5 km)** | | -0.003 | 0.022 |
| **Surface water (10 km)** | | 0.006 | 0.024 |
| **Soil Moisture** | 1 | 0.191*** | 0.031 |
| **Evapotranspiration** | | 0.124*** | 0.012 |
| **Surface water (5 km)** | | 0.008*** | 0.014 |
| **Surface water (10 km)** | | 0.106*** | 0.015 |
| **Soil Moisture** | 2 | 0.357*** | 0.037 |
| **Evapotranspiration** | | 0.074*** | 0.013 |
| **Surface water (5 km)** | | 0.084*** | 0.014 |
| **Surface water (10 km)** | | 0.099*** | 0.015 |
| **Soil Moisture** | 3 | 0.278*** | 0.034 |
| **Evapotranspiration** | | 0.013 | 0.012 |
| **Surface water (5 km)** | | 0.015 | 0.011 |
| **Surface water (10 km)** | | 0.015 | 0.011 |

#### S7.2 WNV MIR models results

Results are reported in the table below, which present coefficient estimates and standard errors (SE) for each hydrological variable and lag. Each row corresponds to a separate regression.

Positive coefficients indicate that higher values of the hydrological variable are associated with higher mosquito infection rates at the specified month lag. Because each hydrological variable is estimated in separate regressions, coefficients should be interpreted as independent associations conditional on fixed effects and covariates. All covariates were standardized.

| Hydrological Variable | Month Lag | Estimate | SE |
|---|---|---|---|
| **Soil Moisture** | 0 | 0.080* | 0.039 |
| **Evapotranspiration** | | 0.05 | 0.010 |
| **Surface water (5 km)** | | -0.009 | 0.011 |
| **Surface water (10 km)** | | 0.008 | 0.011 |
| **Soil Moisture** | 1 | -0.027 | 0.030 |
| **Evapotranspiration** | | -0.005 | 0.009 |
| **Surface water (5 km)** | | -0.023* | 0.009 |
| **Surface water (10 km)** | | -0.016 | 0.010 |
| **Soil Moisture** | 2 | -0.106** | 0.035 |
| **Evapotranspiration** | | 0.014 | 0.010 |
| **Surface water (5 km)** | | 0.011 | 0.007 |
| **Surface water (10 km)** | | 0.007 | 0.010 |
| **Soil Moisture** | 3 | -0.117*** | 0.027 |
| **Evapotranspiration** | | 0.001 | 0.012 |
| **Surface water (5 km)** | | -0.002 | 0.007 |
| **Surface water (10 km)** | | -0.006 | 0.008 |

As a sensitivity check, we look at the effect of standing surface water within the 5- and 10-km buffer of our sampling clusters. In general, results look similar enough between both buffer zones and we go with the 5-km buffer as it may more directly influence mosquito dynamics at our sampling sites.

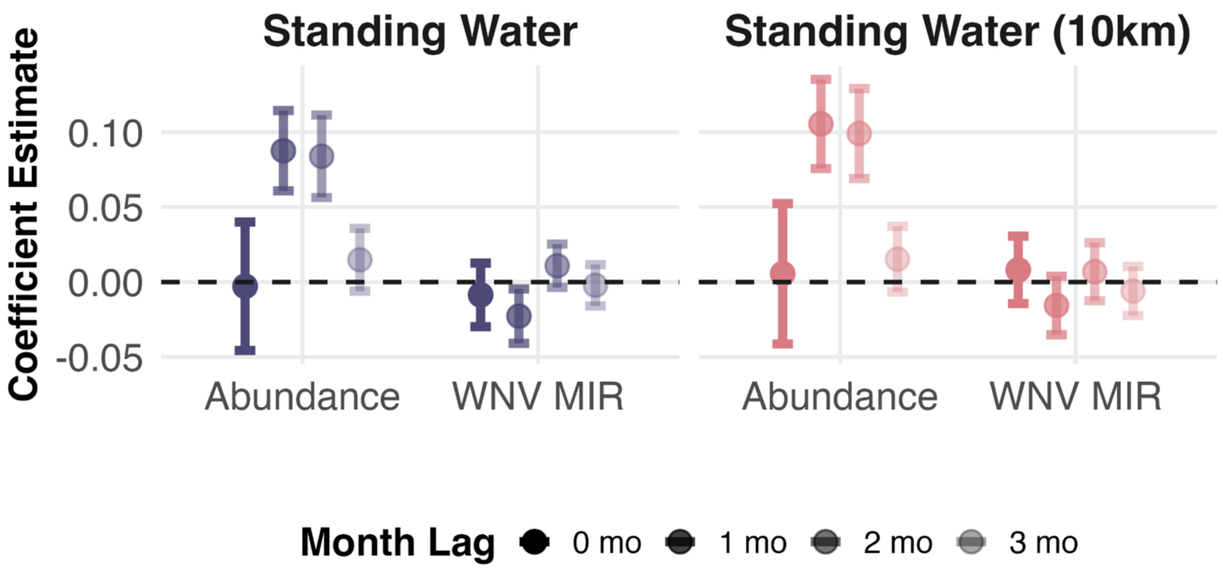

**Text S8 Panel model sensitivity analyses**
To assess the sensitivity of our baseline results, we conducted a series of analyses targeting external validity and measurement uncertainty.

**S8.1 External validity.** We evaluate the generalizability of our findings by re-estimating the baseline model across (i) alternative *Culex* vector species and (ii) additional WNV-endemic regions of California outside the alluvial boundary of the Central Valley. These analyses test whether the estimated drought-mosquito relationship extends across vector taxa (ecological generalizability) and geographic contexts. All models use the Palmer Drought Severity Index (PDSI) as the main effect and include cluster, year and month fixed effects. Standard errors are computed using Conley corrections with a 5 km spatial cutoff.

**S8.1.1 Alternative vector species.** We estimated species-specific models for *Cx. tarsalis*, *Cx. pipiens*, and *Cx. quinquefasciatus*.

Our abundance results show positive associations between PDSI (wetter conditions) and mosquito abundance for *Cx. tarsalis* and *Cx. pipiens*, but not for *Cx. quinquefasciatus*. Effect sizes are largest for *Cx. pipiens*, while estimates for *Cx. quinquefasciatus* are small and not statistically significant.

| **Variables** | ***Cx. tarsalis*** | ***Cx. pipiens*** | ***Cx. quinquefasciatus*** |
|---|---|---|---|
| **PDSI** | 0.153***<br>[0.12, 0.19]<br>(0.0172) | 0.146***<br>[0.10, 0.19]<br>(0.017) | 0.003<br>[-0.06, 0.07]<br>(0.022) |
| **Temp.** | 0.173***<br>[0.08, 0.27]<br>(0.0473) | 0.419***<br>[0.33, 0.51]<br>(0.080) | 0.446***<br>[0.29, 0.60]<br>(0.079) |
| **Temp.$^2$** | -0.218***<br>[-0.26, -0.17]<br>(0.0219) | -0.146***<br>[ -0.19, -0.10]<br>(0.040) | -0.218***<br>[-0.30, -0.14]<br>(0.042) |
| **Obs.** | 82,078 | 48,418 | 36,541 |
| **Clusters** | 1,863 | 1,095 | 877 |
| **$R^2$** | 0.53 | 0.51 | 0.43 |
| **Within $R^2$** | 0.008 | 0.007 | 0.003 |

***p < 0.001; FE, fixed effects. Temp., temperature. Conley standard errors (5 km).

Our West Nile virus minimum infection rate (WNV MIR) results show negative associations between PDSI (drier conditions) and infection rates but is only statistically significant for *Cx. tarsalis*.

| Variables | *Cx. tarsalis* | *Cx. pipiens* | *Cx. quinquefasciatus* |
|---|---|---|---|
| **PDSI** | -0.053***<br>[-0.081, -0.025]<br>(0.0141) | 0.0185<br>[-0.030, 0.025]<br>(0.0213) | -0.0111<br>[ -0.071, 0.049]<br>(0.0307) |
| **Temp.** | 0.087<br>[-0.201, 0.025]<br>(0.0577) | -0.2033***<br>[-0.29, 0.322]<br>(0.0568) | 0.0987<br>[ -0.067, 0.264]<br>(0.0845) |
| **Temp.$^2$** | 0.144***<br>[-0.080, 0.208]<br>(0.0328) | 0.1788***<br>[ -0.092, 0.36]<br>(0.0457) | 0.0205<br>[-0.048, 0.089]<br>(0.0354) |
| **Bird Index** | -0.0045<br>[-0.035, 0.026]<br>(0.0156) | -0.0264<br>[-0.24, 0.024]<br>(0.0153) | 0.0272<br>[-0.012, 0.067]<br>(0.0206) |
| **Obs.** | 25,302 | 19,306 | 13,806 |
| **Clusters** | 1,272 | 727 | 608 |
| **$R^2$** | 0.17 | 0.13 | 0.26 |
| **Within $R^2$** | 0.0029 | 0.0018 | <0.001 |

***p < 0.001. FE, fixed effects. Temp., temperature. Bird, bird community competence index. Conley standard errors (5 km).

Across species, the drought-mosquito relationship is not uniform. For abundance, the pattern broadly aligns with the ecological niches of each vector in the Central Valley. *Cx. tarsalis*, a predominantly rural species associated with agricultural habitats, shows a positive response to wetter conditions. *Cx. pipiens*, which occupies both agricultural (e.g., rice fields in Sacramento Valley) and urban or polluted water bodies, exhibits an even stronger positive association. In contrast, *Cx. quinquefasciatus* - a largely urban species that exploits small water sources such as containers or ornamental vegetation - shows no clear response to PDSI, consistent with weaker dependence on landscape-scale hydrological variability. For infection dynamics, patterns are less consistent, reflecting the greater ecological complexity underlying WNV transmission. While all three species exhibit negative associations between PDSI and infection rates (i.e., higher infections under drier conditions), the effect is statistically significant only for *Cx. tarsalis*. This likely reflects its central role in rural enzootic transmission cycles, where drought-induced concentration of hosts and vectors may amplify transmission. Whereas urban transmission cycles involving *Cx. pipiens* and *Cx. quinquefasciatus* may be buffered by more stable water availability and host resources, weakening the observable relationship with drought.

**S8.1.2 Spatial units outside the alluvial boundary.** To further assess geographic external validity, we re-estimated the baseline model for *Cx. tarsalis* using observations from WNV-endemic regions of California outside the Central Valley alluvial boundary. Specifically, we excluded all observations within the alluvial study domain and instead analyzed data from 1) counties with agricultural presence (Shasta, Riverside) and 2) both agricultural and urban counties (Shasta, Riverside, Orange, and Los Angeles). Because data on bird community competence are not available outside the Central Valley, we restricted this analysis to mosquito abundance models. All specifications are otherwise identical to the baseline, including fixed effects (cluster, year, month) and Conley standard errors (5 km). The estimated effect of PDSI outside the alluvial boundary is highly consistent with the baseline results. In both settings, wetter conditions cause an increase in abundance of *Cx. tarsalis*, and the magnitude of the effect is nearly identical across spatial domains.

| **Variables** | **Within-alluvial boundary** (baseline model) | **Outside boundary** (ag counties) | **Outside boundary** (ag and urban counties) |
|---|---|---|---|
| **PDSI** | 0.153***<br>[0.119, 0.186]<br>(0.017) | 0.065*<br>[0.009, 0.121]<br>(0.029) | 0.0267<br>[-0.024, 0.078]<br>(0.026) |
| **Temp.** | 0.173***<br>[0.080, 0.266]<br>(0.047) | 0.311***<br>[0.154, 0.468]<br>(0.080) | 0.0794<br>[-0.017, 0.176]<br>(0.049) |
| **Temp.$^2$** | -0.218***<br>[-0.260, -0.175]<br>(0.022) | 0.065<br>[-0.005, 0.125]<br>(0.036) | 0.0186<br>[-0.023, 0.060]<br>(0.021) |
| **Obs.** | 82,078 | 20,734 | 33,815 |
| **Clusters** | 1,863 | 329 | 770 |
| **$R^2$** | 0.53 | 0.61 | 0.57 |
| **Within $R^2$** | 0.008 | 0.01 | 0.002 |

***p < 0.001. FE, fixed effects. Temp., temperature. Conley standard errors (5 km).

These results indicate the positive association between wetter conditions and *Cx. tarsalis* abundance is spatially robust in agricultural regions that extend beyond the Central Valley alluvial system. Although the counties included in this analysis differ in landscape composition - ranging from predominantly rural (Shasta) to mixed agricultural-urban (Riverside, Orange) and highly urbanized (Los Angeles) - the consistency of the PDSI effect in agricultural settings suggest the underlying hydrological-ecological mechanism is not confined to a single geographic setting. Notably, while Shasta County lies within the broader Central Valley, it falls outside the defined alluvial boundary used in the main analysis and was therefore excluded from the baseline specification. Its inclusion here, alongside more urbanized counties, reinforces that *Cx. tarsalis* remains responsive to landscape-scale moisture conditions even across heterogeneous environments, provided suitable habitat and temperature conditions are present.

**S8.2 Measurement Uncertainty.** To assess sensitivity to measurement uncertainty in our primary exposure, we replaced PDSI with alternative remotely sensed indicators of hydrological conditions including near-surface soil moisture, standing surface water, and evapotranspiration. Among these, soil moisture exhibited the strongest associations with both mosquito abundance and WNV infection outcomes. However, estimates based on the ESA Climate Change Initiative (CCI; ~ 25 km) had relatively wide confidence intervals, likely reflecting spatial aggregation and measurement error at coarse resolution. We therefore conducted a focused analysis using multiple soil moisture products to evaluate the robustness of results to difference in data sources and spatial scale. CCI was initially used because it provides continuous coverage over the full study period (2003-2023), whereas more recent satellite products such as Soil Moisture Active-Passive (SMAP) are only available from 2015 onward (Reichle et al. 2025). For more contemporary studies post-2015, SMAP is a higher quality data product that measures surface soil conditions at a global scale every two to three days. With this frequency of sampling, soil moisture estimates can better match the collection period of in situ measurements. This would be particularly helpful for studies in more arid regions where water availability is rapidly changing. To balance temporal coverage with measurement precision, we compared results across two widely used datasets CCI (~ 25 km) and SMAP (~ 9 km). These products differ in sensor inputs, retrieval algorithms, and spatial resolution, providing a more rigorous test of sensitivity to measurement error in landscape wetness.

This analysis focuses on *Cx. tarsalis* within the alluvial boundary of the Central Valley, where data availability is most complete. All models retain the baseline specification, including fixed effects (cluster, year, month) and Conley standard errors (5 km). Results from the abundance models are presented below:

| **Variables** | **CCI (~25 km)**<br>(baseline model) | **SMAP (~9 km)** |
|---|---|---|
| **Soil Moisture** | 0.153***<br>[0.119, 0.186]<br>(0.017) | 0.055*<br>[0.0040, 0.11]<br>(0.026) |
| **Temp.** | 0.173***<br>[0.080, 0.266]<br>(0.047) | 0.22*<br>[0.046, 0.40]<br>(0.091) |
| **Temp.$^2$** | -0.218***<br>[-0.260, -0.175]<br>(0.022) | -0.20***<br>[-0.28, -0.11]<br>(0.043) |
| **Obs.** | 82,078 | 690,379 |
| **$R^2$** | 0.53 | 0.52 |
| **Within $R^2$** | 0.008 | 0.012 |

***p < 0.001; *p < 0.05. FE, fixed effects. Temp., temperature. Conley standard errors (5 km).

Results from WNV infection models:

| **Variables** | **CCI (~25 km)** (baseline model) | **SMAP (~9 km)** |
|---|---|---|
| **Soil Moisture** | -0.053*** | -0.010 |
| | [-0.081, -0.025] | [-0.089, 0.068] |
| | (0.0141) | (0.040) |
| **Temp.** | 0.087 | 0.041 |
| | [-0.201, 0.025] | [-0.21, 0.29] |
| | (0.0577) | (0.13) |
| **Temp.$^2$** | 0.144*** | 0.043 |
| | [-0.080, 0.208] | [-0.057, 0.14] |
| | (0.0328) | (0.051) |
| **Bird Index** | -0.004 | -0.069** |
| | [-0.04, 0.21] | [-0.12, -0.018] |
| | (0.016) | (0.026) |
| **Obs.** | 25,302 | 12,516 |
| **$R^2$** | 0.17 | 0.20 |
| **Within $R^2$** | 0.0029 | 0.0014 |

***p < 0.001. FE, fixed effects. Temp., temperature. Conley standard errors (5 km).
